# Cruciform-forming AT/TA repeats are acted upon by structure selective endonucleases and Rad51 prior to repositioning to the nuclear periphery for repair

**DOI:** 10.64898/2026.01.20.700634

**Authors:** Jan Leendert Boer, Sara M. Crippen, Amelia J. Kim, Daphne N. Kramer, Bertrand Theulot, Daniel Dovrat, Amir Aharoni, Duncan J. Smith, Catherine H. Freudenreich

## Abstract

Structure-forming DNA repeats can pose a barrier to DNA replication and repair, creating chromosomal fragile sites. An AT/TA DNA repeat, derived from the Flex1 region of human common fragile site FRA16D, can form hairpin and cruciform structures which interfere with DNA replication. When inserted into the *S. cerevisiae* genome, the Flex1(AT)_34_ repeat stimulates chromosome deletions in a manner dependent on the Mus81-Mms4 nuclease and the SLX4 nuclease scaffold. It was previously found that hairpin forming CAG/CTG repeats move to the nuclear periphery to maintain genomic stability. Here, we show that a structure forming AT/TA repeat also relocalizes to the nuclear periphery in late S/G2 phase in a replication and length-dependent manner. In contrast to the CAG repeat, this shift in nuclear positioning is dependent on polySUMOylation and the activity of the Mus81-Mms4 nuclease. Processing by the Mre11 nuclease and Rad51-dependent strand exchange occurs prior to repositioning. Replication analysis indicates that the replisome likely bypasses the AT/TA repeat leaving behind a DNA structure that initiates relocation to the nuclear periphery. We conclude that AT/TA repeats form post-replicative DNA structures that are targeted for nuclease cleavage and require hairpin processing and repositioning to the nuclear periphery for homologous recombination-dependent repair.

## Introduction

Secondary structures formed by DNA repeats pose a significant barrier to replication and repair and are frequent sites of genome instability (Poggi and Richard 2021; Brown and Freudenreich 2021; Wang and Vasquez 2023; Boer et al. 2025). One noteworthy sequence that forms hairpin and cruciform structures are AT/TA dinucleotide repeats (AT repeats) (Kaushal and Freudenreich 2019). How AT repeats are faithfully replicated and repaired despite non-B DNA structure formation remains incompletely understood.

Common fragile sites (CFS) are genomic regions prone to breakage, deletions and translocations. In human cells under replication stress these loci do not finish replication in S phase and present as gaps and breaks in metaphase spreads. Under-replication of CFSs and resulting fragility is thought to be influenced by a variety of characteristics including transcription of large genes, delayed replication, spanning of TAD boundaries, and AT content (Sinai and Kerem 2023). One of the most frequently occurring CFSs in the human genome is FRA16D, which is located in the WWOX (WW domain-containing oxidoreductase) gene, a tumor suppressor gene (Taouis et al. 2021). This fragile site contains six AT-rich subregions termed Flex1-6 (Ried et al. 2000). Flex1 contains a polymorphic AT/TA rich repeat that ranges from 11 to 88 repeats in humans (Finnis et al. 2005). AT repeats above 22 repeat units are known to form a cruciform secondary structure *in vivo* (McClellan et al. 1990; Dayn et al. 1991; Cote and Lewis 2008; Kaushal et al. 2019) and long AT repeats have been shown to cause chromosome fragility in a variety of systems and species (Sinai and Kerem 2023). Indeed, structure-forming AT-rich repeats are the most common sites of chromosome breakage retrieved in human cells that have experienced replication stress (Shastri et al. 2018). Insertion of the Flex1 region into a yeast chromosome led to an increase in chromosome end loss or chromosome deletions in an AT repeat-length dependent manner compared to insertion of other regions of the FRA16D locus that are not predicted to be structure forming (Zhang and Freudenreich 2007; Kaushal et al. 2019). Upon aphidicolin (APH) or hydroxyurea (HU) induced replication stress, the Flex1 AT repeat stimulated increased homologous recombination compared to a control sequence in both yeast and human U-2OS cell lines (Wang et al. 2014).

The Flex1(AT)_34_ repeat was shown to stall replication *in vivo* in yeast cells by two-dimensional (2D) gel electrophoresis (Zhang and Freudenreich 2007), and human polymerase delta (Pol δ) has difficulty replicating through the AT tract *in vitro* (Kaushal et al. 2019). A recent study showed that translesion polymerase zeta (Pol ζ) plays a role in preventing AT repeat-mediated chromosome breaks (Das et al. 2024). *In vitro,* Pol ε is severely stalled at the AT hairpin base whereas both human and yeast Pol ζ can enter the hairpin base and replicate through the AT repeat in a stepwise manner (Das et al. 2024).

Several lines of evidence implicate structure selective endonucleases (SSEs) in targeting cruciform-forming AT/TA repeats to cause DNA breaks. Using a yeast plasmid based system, the yeast Mus81-Mms4 SSE was shown to cleave a cruciform structure formed by an AT/TA repeat from the human neurofibromin 1 locus (Cote and Lewis 2008). The SSEs associated with the Slx4 scaffold protein, including Mus81-Mms4, Slx1, and Rad1-Rad10, were required for chromosome end loss or internal deletions stimulated by cruciform-forming lengths of the Flex1 AT repeat from the human FRA16D locus integrated at a yeast chromosomal site (Kaushal et al. 2019). In cancer cell lines that have accumulated expanded AT/TA repeat tracts due to microsatellite instability, depletion of WRN led to 2-ended double-stranded breaks (DSBs) at long AT repeats that were dependent on the human MUS81-EME1 nuclease (van Wietmarschen et al. 2020; Matos-Rodrigues et al. 2022). In the yeast system, it was shown that the cleavage of the Flex1-derived (AT)_34_ repeat structure resulted in hard to repair intermediates due to the formation of secondary structures within and flanking the AT repeat that prevented efficient healing of the break (Kaushal et al. 2019). Thus, the observed fragility of regions containing structure-forming repeats are likely a combination of their propensity to be cleaved by SSEs and their subsequent difficulty to repair due to hairpin-capped ends (Kaushal and Freudenreich 2019; Cejka and Symington 2021; Lobachev, Gordenin, and Resnick 2002).

The replication of repetitive DNA elements occurs in the context of chromatin and the spatial organization of chromosomes in the nucleus. Certain types of DNA damage reposition to the nuclear periphery (NP) for repair and associate with the nuclear pore complex (NPC). These include persistent DSBs, DSBs in heterochromatin, eroded telomeres, and some types of replication impediments (reviewed in (Boer et al. 2025; Schirmeisen, Lambert, and Kramarz 2021; Gasser and Stutz 2023; Chiolo et al. 2025). This mechanism appears to be conserved, as DSBs induced by topoisomerase poison etoposide or by nucleases in transcriptionally active regions associate with the NP in human cells (Shokrollahi et al. 2024; Le Bozec et al. 2024). In terms of replication barriers, long tracts of CAG repeats that form DNA secondary structures were shown to relocalize to the NPC in a manner dependent on their replication (Su et al. 2015). G4 forming telomeric sequences move to the NPC from a clustered nuclear envelope position upon replicative senescence (Khadaroo et al. 2009; Churikov et al. 2016) and stalls in telomeric DNA that occur upon replication stress cause repositioning to the NPC (Aguilera et al. 2020; Pinzaru et al. 2020). A protein mediated replication fork stall in *S. pombe* repositions to the NPC, and this process facilitates Pol δ-mediated fork restart (Kramarz et al. 2020; Schirmeisen et al. 2024). Aphidicolin treatment, which stalls or collapses replication forks, caused a decrease in the distance of replication foci to the NP in human U-2OS cells (Lamm et al. 2020), and NPC components were found enriched at replication forks stalled for 2 hours with 5μM APH in immortalized mouse embryonic fibroblasts (Rivard et al. 2024). However, not all types of replication barriers provoke relocation to the NP. In yeast, levels of hydroxyurea (HU) that transiently stall replication or base damage by methyl-methane sulfinate (MMS) do not have this effect, but longer treatments of HU or HU+MMS that cause fork collapse do (Nagai et al. 2008).

Based on the knowledge that forks stalled by long hairpin-forming tracts of CAG/CTG repeats and replication stress within G4-forming telomeric sequences cause repositioning to the NPC to maintain genome stability, we asked whether the structure-forming Flex1(AT)_34_ repeat also provokes repositioning to the NPC. In this study, we show that a cruciform forming AT/TA repeat relocalizes to the NPC in late S/G2 phase in a replication and length-dependent manner. The cleavage of structure selective endonucleases, including Mus81-Mms4, Slx1-Slx4, Rad1-Rad10, and Sae2-Mre11 is a critical requirement for repositioning, and Mus81-Mms4 cleavage products are bound by Rad51 prior to movement to the NPC. The repositioning also depends on polySUMOylation by Nse2/Mms21, Siz1, and Siz2 SUMO ligases, and SUMO-targeted ubiquitin ligases Slx5-Slx8 and Uls1. Interrupting this relocation pathway causes an increase in break-stimulated recombination, resulting in deletion of the repeat tract and flanking sequences. Despite the replication dependence of relocation, an (AT/TA)_34_ repeat does not lead to a significant change in the chromosomal replication profile, suggesting that the barrier is bypassed. We propose a model whereby unreplicated cruciform structures that are left behind after passage of the replication fork are targeted for nuclease cleavage, which initiates Rad51-dependent strand exchange and repositioning to the NPC for repair.

## Methods

### Yeast strains and genetic manipulation

*S. cerevisiae* strains used for microscopy had the Flex1 (AT/TA)_n_ repeat of varying sizes or a Flex1 control sequence (no repeat control), a ∼380 bp sequence from human FRA16D that is AT-rich but not predicted to form any secondary structure (Zhang and Freudenreich 2007; Kaushal et al. 2019), inserted into a yeast chromosome VI noncoding region upstream of the tA(AGC)F gene. The location is 6051 bp downstream from ARS607. A *lac* operator array was present 6,452 bp from the start of the repeat or the control sequence in the GFP-LacI GFP-Nup49 strain and 6,431 bp away from the LacO array in CFP-LacI mCherry-Nup49 strains. The parent Flex1(AT)_34_ and the control sequence Direct Duplication Recombination Assay (DDRA) strains were previously constructed and from Kaushal et al. (2019). Mutant strains were constructed by one-step gene replacement and verified by a combination of PCR sizing and sequencing. AT repeat length was verified in the wild-type (WT) strain by sequencing and checked in derived strains by PCR or sequencing. For *S. cerevisiae* strains used for replication fork progression experiments, the AT repeat or control inserts were placed between two arrays ∼22 kb apart on chromosome IV, 128xlacO bound by LacI-Envy and 128xtetO bound by TetR-tdTomato, a location replicated primarily by ARS413 (Dovrat et al. 2018). The start of the lacO array is ∼3 kb downstream from ARS413, the start of the inserts is ∼12 kb downstream from ARS413, and the start of the tetO array is ∼31 kb downstream from ARS413. Yeast knock-in insert fragments were made using either PCR or by digestion of a plasmid which contains the insert and homology up and downstream of the target locus, transformed into yeast cells along with a Cas9 plasmid with a guide targeting NATMX, resulting in replacement of NATMX with the desired insert. All yeast strains are listed in Table S1.

### Microscopy

Yeast were grown overnight in yeast complete (YC) media and diluted back to an optical density OD_600_ of ∼0.1. After 2-4 doublings in YC media at 30°C, cells were prepared for imaging by allowing the yeast to settle for 10 minutes on Ibidi uncoated 8 well slides coated with concanavalin A. Cells were fixed using a 4% final concentration solution formaldehyde and washed 3x using YC or phosphate buffered saline (PBS). Alternatively, cells were fixed with 4% formaldehyde and washed 3x using PBS in a 1.5mL microcentrifuge tube, and agar pads were prepared on depression slides using 1.4% YC agar. 5μl of cells were placed on the agar pad followed by a coverslip.

Z-stack images were taken using a Delta Vision Ultra High-Resolution microscope at 100x magnification. Slices were 0.2μm and 25 Z-planes were taken per field of cells. Exposure time for microscopy experiments were 100ms for GFP, 100ms for mCherry, 100ms for CFP, 100ms for i-mVenus, and 50ms for brightfield. Images were deconvolved and zoning was scored in mid to late S phase cells with the criteria of the bud being at least 20% the size of the mother cell and having a round nucleus except for experiments done with synchronization. The ImageJ point picker program was used to quantify zoning as described previously in Su et al. (2015).

For live cell colocalization timing experiments cells were arrested in α-factor and allowed to settle on Ibidi uncoated 8 well slide coated with concanavalin A for the last 10 minutes of the arrest. Cells were then washed 5x with prewarmed YC media and imaged every 5 minutes. For fixed cell colocalization experiments, after cell cycle arrest with α-factor, cells were washed 3x with prewarmed YC and allowed to grow in YC media while shaking at 30°C. Timepoints were taking 10 minutes prior to the indicated time, allowed to settle for 10 minutes at 30°C and then arrested with 4% final concentration solution formaldehyde. Images were then taken as described above. For co-localization analysis, overlapping or touching mCherry-Nup49 and LacI-CFP foci were scored as colocalized while non touching foci were deemed not colocalized.

### Okazaki Fragment Sequencing

Cells carrying a doxycycline-repressible *CDC9* gene were grown to mid-log phase before addition of 40 mg/L doxycycline for 2 hrs. Genomic DNA was purified from spheroplasts and Okazaki fragments were enriched, purified and prepared for Illumina sequencing as previously described Smith and Whitehouse (2012). Paired-end sequencing was carried out on a NextSeq 500. After sequencing, read mapping was performed with BWA-MEM (v0.7.17) on the *Saccharomyces cerevisiae* S288C reference genome (vR64-5-1-20240529). Using Samtools (v1.19), reads with multiple alignments (SAM flag 0×100) and supplementary alignments (0×800) were filtered out. Strand-specific coverage was extracted using Samtools flags. For the reverse strand, reads with 0×50 (read 1, reverse complement) and 0×90 (read 2, reverse complement) were selected. For the plus strand, reads with 0×40 (read 1, forward) and 0×80 (read 2, forward) were extracted. Coverage was computed using Samtools depth with the -aa flag to include all positions, even those with zero reads aligned. Coverage files for both strands were merged using awk. Coverage profiles and fork directionality profiles were plotted using a custom R script.

### Direct Duplication Recombination Assay

Yeast strains were first patched onto YC-Ura plates to ensure the starting patches had the Flex1(AT)_34_ repeat and the *URA3* gene. From this patch the yeast were plated for single colonies on yeast extract peptone dextrose (YEPD) plates. The colonies were allowed to grow without selection for three days at 30°C. Following this growth, 10 colonies were randomly selected and half the colony was resuspended in 400μl of diH_2_O. From this solution a 1:400 dilution was plated on 5-Fluoroorotic acid (FOA) -Ade plates to select for loss of the Flex1(AT)_34_ *URA3* cassette and reconstitution of the *ADE2* gene. A portion of each colony resuspension was combined, and dilutions of 10^-4^ and 10^-5^ were plated on YEPD plates to obtain a total cell count. Using the colony counts from the 5-FOA -Ade and YEPD plates and utilizing the method of the median, a rate of 5-FOA resistance was calculated (Lea and Coulson 1949).

### Rad51 and Rfa3 colocalization experiments

A functional internally tagged Rad51-i-mVenus gene was created by replacing GFP with mVenus in plasmid pAT624 (Rad51-GFP plasmid), a gift from Angela Taddei, (Liu, Mine-Hattab, et al. 2023). This was done using the NEB HiFi assembly kit, to create pCF913 containing an internally tagged Rad51 construct; as in the original Rad51-i-GFP construct, the mVenus tag is inserted 162 bp (21 amino acids) from the Rad51 start codon and is flanked by 48 bp linkers. The knock in fragment was created from this plasmid by digestion with KpnI-HF and HindIII-HF. The knock-in fragment was transformed into yeast cells along with pAT569, a Cas9 plasmid with a guide targeting Rad51(Liu, Mine-Hattab, et al. 2023). Recombinants were verified by PCR sizing, sequencing, and microscopy.

A C-terminally tagged Rfa3-mVenus protein was created by PCR amplifying a Rfa3-mVenus gene adjacent to a KanMX marker from a previously constructed yeast strain. The knock-in was verified by PCR sizing, sequencing, and microscopy.

For Rad51 or Rfa3 colocalization experiments, microscopy slides were prepared as described above. Mid to late S phase cells were counted as having a round nucleus and a bud being at least 20% the size of the mother cell. The distance between Rad51 or Rfa3 foci and the LacO array that marked the approximate location of the Flex1(AT)_34_ or no repeat control sequence locus, nucleus diameter and the distance from the NP, marked by mCherry-Nup49, to the LacO-LacI-CFP focus was measured. A distance between the mVenus focus and the LacO/LacI-CFP focus of under 0.3μm was used as the cut off for colocalization. The ratio of distance to the NP over the radius of the nucleus was used to determine the zone.

### Live cell microscopy for monitoring replication fork progression

Cells were grown overnight in YC media and diluted back to an OD_600_ of ∼0.1. Cells were allowed to grow to an OD_600_ of ∼0.4-0.6 followed by arrest for 2 hours with 10 μg/ml alpha-factor. In the last 10 minutes of arrest 400 μl cells were vigorously vortexed and allowed to settle on an Ibidi 8 well slides coated with concanavalin A at 30°C. Cells were then washed 5x with prewarmed YC media to release them into S phase and immediately imaged in the microscope chamber at 30°C for the duration of the experiment. Ten fields with cells were identified and imaged in each experiment. Live-cell imaging using a DeltaVision microscope was performed at 1 min intervals for a 3 h period. Images were taken with the 60x oil objective and the B-G-O-FR polychroic filter. Channel parameters were polarized light at 50 ms, orange (542 nm) for 20 ms and green (475 nm) at 20 ms. Twelve 0.5 μm slices were taken for each of the 10 fields.

### Quantification and statistical analysis

Graphs and statistical analysis were performed using Prism 10 software. For relocalization or colocalization experiments statistical significance was calculated using Fisher’s exact test. For DDRAs a t-test with Welch’s correction was employed. For live-cell microscopy data quantification, images were analyzed with a custom made computational pipeline based on Python (AutoCRAT) to measure replication fork progression, essentially as previously described Dovrat et al. (2018). AutoCRAT allows the identification, tracking, and quantification of fluorescence signals from LacI-Envy and TetR-tdTomato, foci in single cells. For each strain we measured an average of 320 cells (range 185-679 except for the *rev3* no repeat control at 33). Independent experiments for each strain are shown in Figure S6B. Statistical analysis of replication times was analyzed using Monte Carlo resampling with 1,000,000 iterations.

## Results

### The Flex1(AT)_n_ repeat localizes to the nuclear periphery in late S/G2 phase in a length and replication dependent manner

To assess if a structure-forming AT repeat relocalizes to the nuclear periphery, a zoning assay was employed as described in Su et al. (2015). The Flex1 sequence from the human FRA16D CFS containing various lengths of an AT repeat, or an AT-rich control sequence from FRA16D not predicted to form a DNA structure termed the no repeat control, were inserted downstream of ARS607 on *Saccharomyces cerevisiae* chromosome VI. This locus was visualized by a LacO array bound by GFP-LacI 6.4 kb upstream of the repeat. Nup49 was N-terminally labelled with GFP or RFP, allowing for visualization of the NP (Figure 1A). The nucleus was split into 3 equal zones with zone 1 being the area closest to the NP (Figure 1A). Cells in mid to late S phase, as determined by a round nuclear morphology and cell bud size, were analyzed. Previous data from our lab showed that a (CAG/CTG)_130_ repeat localizes to the NPC in a length dependent manner, with the repeat locus maximally found at the NPC 60 minutes into S phase, in ∼50% of cells (Su et al. 2015; Whalen et al. 2020). Compared to the control sequence the (AT)_14_ repeat is not significantly enriched in zone 1. However, loci with repeat tract lengths equal to or over 23 are enriched in zone 1 compared to the control sequence, with the Flex1(AT)_34_ sequence found in zone 1 62% of the time (Figure 1B; Figure S1A). In Kaushal et al. (2019), using the same sequences as this study, a S1 nuclease cleavage assay showed that the (AT)_23_ and (AT)_34_ repeats are able to form a cruciform structure while the (AT)_14_ and the control sequences do not. This suggests that formation of the AT/TA cruciform structure contributes to the repositioning to the NP.

**Figure 1:**
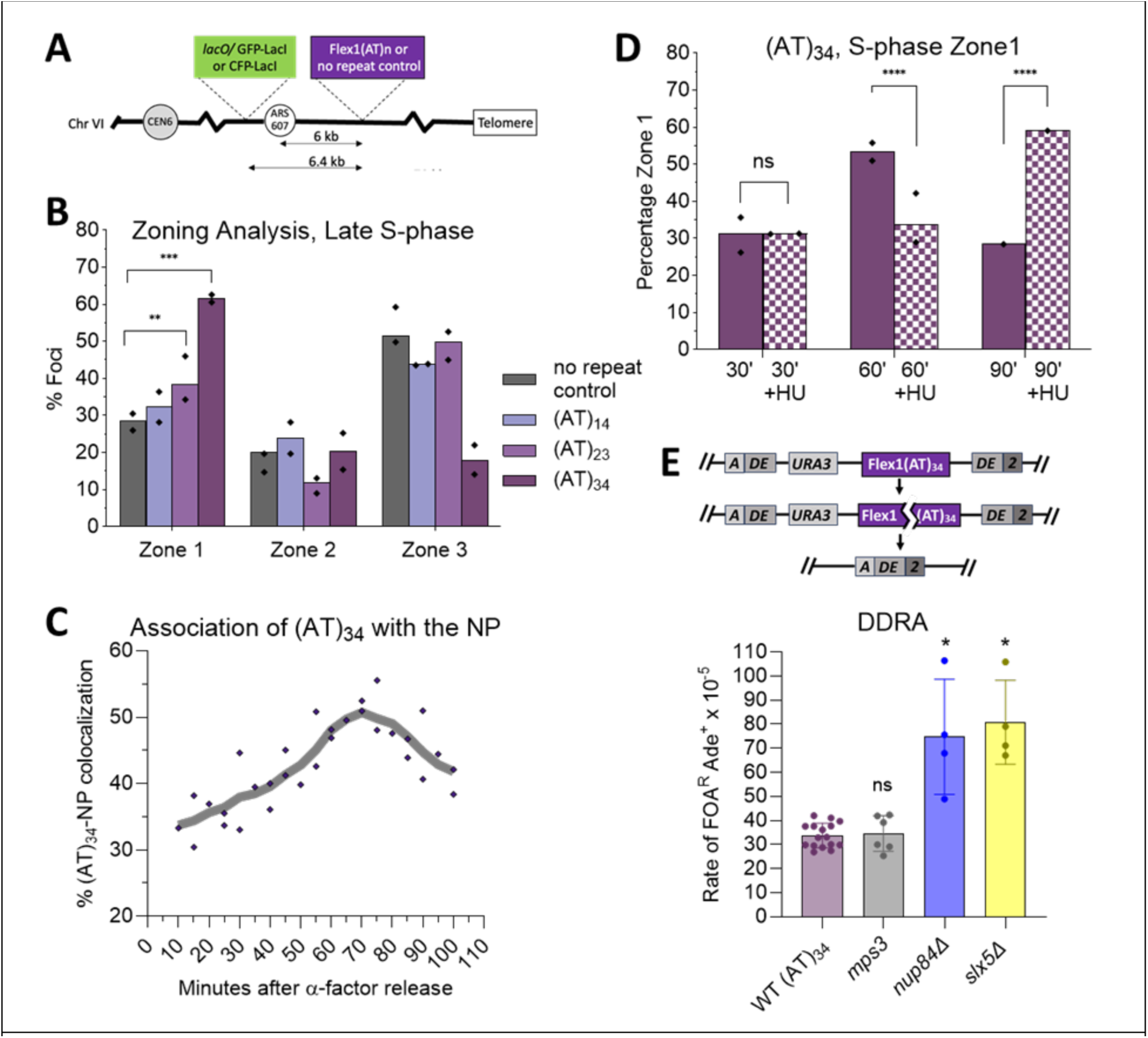
The structure forming AT/TA repeat localizes to the nuclear periphery in late S/G2 phase in a length and replication dependent manner. A) Yeast Chromosome VI with a *LacO* array inserted 0.3 kb upstream of ARS607. The Flex1(AT)_n_ repeat sequence or the AT-rich no repeat control sequence from FRA16D were inserted 6051 bp downstream of ARS607. The locus is visualized with LacI-GFP or LacI-CFP. The NPC is visualized by GFP-Nup49 or RFP-Nup49. Z stack images of the nucleus were divided into 3 zones by area as depicted and the focus assigned to a zone as described in the methods. B) The percentage of GFP foci in each zone in cells identified as being in mid to late S phase by bud size for various (AT)_n_ repeat lengths and the control sequence; the percent of all cells counted is plotted (n>190), points indicate the mean values from 2 independent isolates (n>91 each). C) Percentage of colocalization of the LacI-CFP/LacO focus marking the (AT)_34_ locus with the NP marked by m-Cherry tagged Nup49 at the indicated time points after alpha factor arrest in G1 and release into S phase. Individual data points from 1 live cell experiment and 2 fixed cell experiments are shown (plotted individually in Fig. S1C); the line is a mean with 2^nd^ order smoothing (4 neighbors). D) Zoning assay of cells after alpha factor arrest and release into YC or YC+0.2M HU containing media, done as in B. E) Rate of (AT)_34_-stimulated deletions as measured by a direct duplication recombination assay (DDRA); points indicate independent experiments. For zoning assays, pairwise statistical comparisons by Fisher’s exact test; **p<0.01, ***p<0.001, ****p<0.0001. Zoning data n values are between 100-400 cells per strain or condition; 2 independent isolates were tested for each strain, indicated by data points on bars; exact n values, zone 1 percentages and p values are listed in Table S2 and S3. Colocalization data n values are between 99 and 189 cells per timepoint; exact n and colocalization values for each experiment are in Table S4. For DDRAs, statistical comparison by Welch’s t-test, *p<0.05. Each strain tested with 2 independent isolates, rates and p values are listed in Table S5.

To investigate the exact timing in S phase of the Flex1(AT)_34_ locus repositioning to the NP, cultures were arrested in α-factor, which halts the cell cycle at the G1/S boundary, and released. Samples were analyzed at indicated time points post release. When analyzed by zone, the percentage of cells with the (AT)_34_ locus in zone 1 increases gradually over the cell cycle to a maximum of 55% at 75 minutes after G1 release but sharply drops by the 85 minute time point (Figure S1B). Since the zoning assay only analyzes cells with round nuclei typical of S phase cells, this indicates that cells have progressed out of S phase by 85 minutes. Right before mitosis the yeast nucleus moves to the bud neck and is no longer round, making a zoning assay not an appropriate way to ascertain the distance to the NP in G2 phase cells. To more accurately gauge association of the (AT)_34_ locus with the NP at these late S/G2 timepoints, a colocalization assay was employed in which overlap between the marked chromosomal locus and the NP was scored. The (AT)_34_ repeat locus is gradually enriched at the NP starting at 40-50 minutes post G1 release (Figure 1C; Figure S1C). Maximal (AT)_34_ relocalization occurs at 60-75 minutes post α-factor release, after which the NP association decreases. Most cells entered G2 phase between 65-75 minutes after α-factor release as determined by nuclear morphology. This shows that the (AT)_34_ locus is maximally associated with the NP in late S/G2 phase.

To investigate if replication fork progression through the (AT)_34_ repeat is necessary for the relocalization to the NP cells were treated with hydroxyurea (HU), which inhibits replication by decreasing dNTP levels (Schilsky et al. 1992) and oxidizing replicative DNA polymerases, causing their dissociation (Shaw et al. 2024). Cells were released from G1 arrest into media containing 0.2M HU, which stalls replication forks ∼3-5 kb from the origin (Cobb et al. 2003) so that most replication forks will not reach the (AT)_34_ repeat which is located ∼6 kb away. For short treatments of 1 hour or less, the replication fork does not collapse and can be restarted upon removal of the HU (Petermann et al. 2010). For prolonged HU treatments of 2 hours, the replication fork can collapse and reposition to the nuclear periphery (Nagai et al. 2008). At 30 minutes after α-factor release no enrichment at the NP occurs in either condition (Figure 1D). This is consistent with the data from the live cell time course, as little repositioning is observed at that time point (Figure 1C). At 60 minutes post release, the yeast released into synthetic media exhibit ∼55% zone 1 occupancy while the yeast released into media containing 0.2M HU are not enriched in zone 1 (Figure 1D). This reveals that replication fork progression through the (AT)_34_ repeat is necessary for repositioning and suggests that repositioning. At 90 minutes post release, the tracked region is no longer enriched in zone 1 in the no drug condition while HU treated cells show enriched zone 1 occupancy (Figure 1D). This could be due to fork collapse occurring by this timepoint or due to the slowed fork finally reaching the AT repeat tract.

Most DNA damage studied to date that relocates to the nuclear periphery interacts with components of the nuclear pore (Boer et al. 2025). In *S. cerevisiae*, persistent DSBs and expanded CAG repeats both interact with Nup84 of the Y complex through the Slx5 protein, and HO-induced DSBs also interact with the SUN domain protein Mps3 embedded in the inner nuclear membrane (Nagai et al. 2008; Su et al. 2015). To gain insight into the role of AT repeat anchoring at the nuclear periphery the consequences of removing Nup84 or Mps3 function on repair of repeat-induced breaks were tested. A Direct Duplication Recombination Assay (DDRA), an assay that measures single strand annealing (SSA) events, a proxy for chromosome breaks, was caried out in *nup84Δ*, *slx5Δ*, and *mps3Δ75-150* mutants. Previously, insertion of the Flex1(AT)_34_ repeat was shown to increase the rate of SSA compared to a no repeat control sequence in a length dependent manner (Kaushal et al. 2019). An increase in the rate of Flex1(AT)_34_-stimulated deletions was seen in both *nup84Δ* and *slx5Δ* mutants compared to the WT but not in a *mps3Δ75-150* mutant (Figure 1E). This indicates that Nup84, Slx5 and NPC anchoring are important in maintaining the (AT)_34_ repeat to prevent deletions.

Taken together, these results show that the structure forming AT/TA repeat relocates to the NPC in a length and replication dependent manner in late S/G2 phase, and that disruption of this pathway leads to more deletions of the repeat locus. The longer the AT/TA repeat, the greater the percentage of cells that exhibited locus repositioning to the nuclear periphery.

### Structure selective endonucleases are required for the repositioning of the (AT)_34_ repeat to the nuclear periphery

Previously, all three structure-specific nucleases associated with the Slx1-Slx4, Mus81-Mms4, Rad1-Rad10 (SMR) super complex were shown to be required for Flex1(AT)_34_ repeat-mediated deletions (Kaushal et al. 2019). These results implied that the SMR-associated nucleases were either required for causing a break that stimulated SSA, or for facilitating healing by SSA, or both. The effect of the Mus81-Mms4 nuclease was specific to structure-forming AT repeat lengths and had no effect on the no repeat control sequence (Kaushal et al. 2019). To test if the SMR complex is necessary for AT repeat-mediated repositioning, zoning analysis was performed on mutants. All the tested components of the SMR complex exhibited a decrease in the frequency that the repeat locus is detected in zone 1 (Figure 2A; Figure S2A). Mus81 and its obligate binding partner Mms4 (Fu and Xiao 2003) as well as the SLX4 scaffold and its associated nuclease Slx1 all had similar and statistically significant decreases. Deletion of Rad1 of the Rad1/Rad10 nuclease which can cleave at the base of hairpin loops (Lu et al. 2015) was also required. This result is in stark contrast to the (CAG)_130_ repeat where a deletion of Mus81 does not change the enrichment of the (CAG)_130_ repeat at the NP in mid to late S phase (Figure 2B). To determine whether this requirement was specific to the SMR complex nucleases, Yen1, a SSE that is not part of the SMR complex (Wild and Matos 2016) was tested. This mutant does not show a decrease but interestingly shows a slight increase in zone 1 repeat enrichment (Figure 2A). Evaluation of the timing of relocation in the *yen1Δ* strain did not reveal a notable difference from the WT (Figure S2B). We conclude that cleavage by the SMR complex is a required step in the repositioning of the (AT)_34_ repeat to the NPC.

**Figure 2:**
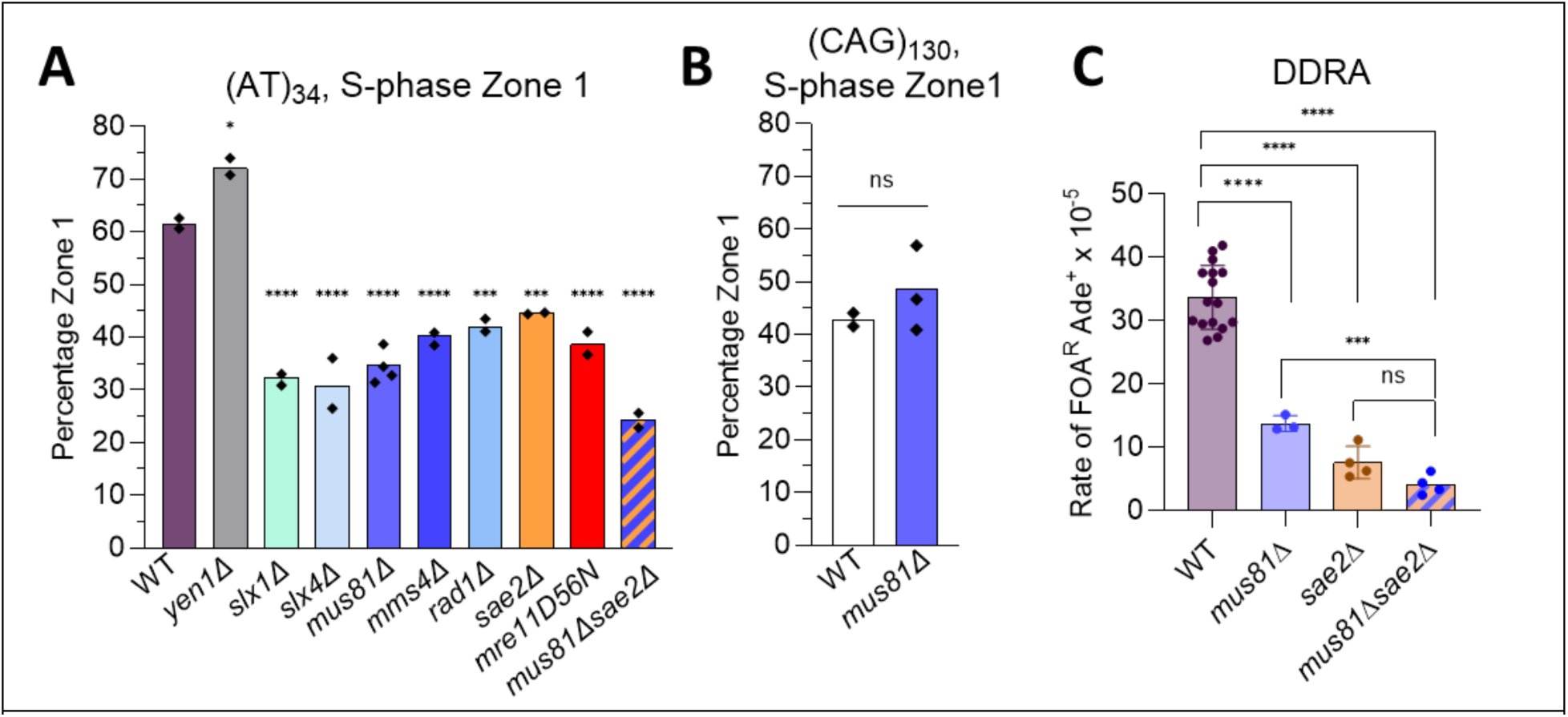
Deletion of structure selective endonucleases reduces zone 1 occupancy of the Flex1(AT)_34_ repeat. A) Rate of (AT)_34_-stimulated deletions in SSE mutants as measured by the DDRA; *mus81Δ and sae2Δ* data from (Kaushal et al. 2019). B) Percentage GFP foci in zone 1 during mid to late S phase for SSE mutants containing Flex1(AT)_34_ integrated on chr VI, or C) containing a (CAG/CTG)_130_ tract at the same locus, WT (CAG)_130_ data from (Whalen et al. 2020). For zoning assays, pairwise statistical comparisons by Fisher’s exact test; *p<0.05, ***p<0.001, ****p<0.0001. Zoning data n values are between 100-400 cells per strain; 2 independent isolates were tested for each strain with percents obtained for each indicated by data points on bars. Exact n values, zone 1 percentages and p values are listed in Table S3. For DDRAs, points indicate independent experiments. pairwise statistical comparison by Welch’s t-test; ***p<0.001, ****p<0.0001. Each strain tested with 2 independent isolates; rates and p values are listed in Table S5.

The AT-rich nature of Flex1(AT)_34_ could cause the formation of hairpin capped ends upon DSB induction by the SMR complex which can inhibit healing of the break. Sae2 stimulates the Mre11-Rad50-Xrs2 (MRX) complex to cleave hairpin capped ends (Tamai et al. 2024) and Sae2 was previously implicated in repair of the cleaved Flex1(AT)_34_ sequence as a *sae2Δ* mutant showed a specific decrease in AT repeat-stimulated deletions (Kaushal et al. 2019). Deletion of Sae2 led to a significant a decrease in zone 1 enrichment during mid to late S phase. Furthermore, a Mre11 nuclease dead mutant, *Mre11D56N* (Moreau, Ferguson, and Symington 1999), exhibited a similar decrease in zone 1 occupancy (Figure 2A). These results suggest a role for hairpin processing in repositioning of the repeat to the periphery. To determine if both cleavage by Mus81 *and* processing of a potential hairpin capped ends by Sae2-Mre11 are necessary for optimal repositioning, Mus81 was knocked out in conjunction with Sae2. This double mutant showed a decrease in zone 1 occupancy down to ∼20% (Figure 2A) showing a potential sequential role for both nucleases. To see if this additive effect can also be observed in the DDRA, a double *mus81Δsae2Δ* mutant was created. Comparing the SSA rate of this strain with previously published single mutants (Kaushal et al. 2019), the double mutant shows a statistically significant decrease compared to the *mus81Δ* single mutant revealing a potential additive effect of these two mutants (Figure 2C). There is no statistically significant difference between the DDRA rate in a *rad52Δ* strain (Table S5), required for SSA, and *mus81Δsae2Δ* mutants indicating that the lower limit of SSA in the assay has been reached, illustrating the critical role being played independently by Mus81 and Sae2.

Altogether, the analysis of nuclease mutants suggests that SSEs are acting on the structure that is formed by the (AT)_34_ repeat and that both the SMR complex and Sae2-stimulated Mre11 nuclease activity play a crucial role in allowing repositioning of the repeat to the NPC. The fact that *mus81Δsae2Δ* double mutants showed an additive effect in both the zoning and fragility assays suggests that Mus81-Mms4 and Mre11 nuclease activities have separate but complementary roles in this pathway.

### Relocalization of the (AT)_34_ repeat is dependent on polySUMOylation and SUMO targeted Ubiquitin Ligases Slx5/8 and Uls1

Relocalization of a (CAG)_130_ repeat to the NPC is dependent on SUMOylation of target proteins, including RPA, Rad52 and Rad59 by the SUMO E3 ligase Mms21 (hNSE2). To test whether SUMOylation plays a role at the (AT)_34_ repeat, a *smt3-331* mutant of the yeast SUMO protein Smt3, which reduces overall Smt3 SUMOylation levels, was tested. This strain had a significant decrease in zone 1 positioning (Figure 3A). A more specific *smt3-KallR* mutant was then tested, in which all lysine residues are mutated to arginine residues allowing only monoSUMOylation to occur (Bylebyl, Belichenko, and Johnson 2003). The *smt3-KallR* mutant showed a similar and significant decrease in zone 1 occupancy to 23% (Figure 3A), which is at or below the baseline no repeat control level of 29% (Figure 1B). This result indicates that monoSUMOylation is not sufficient for anchoring the AT repeat damage at the NPC.

**Figure 3:**
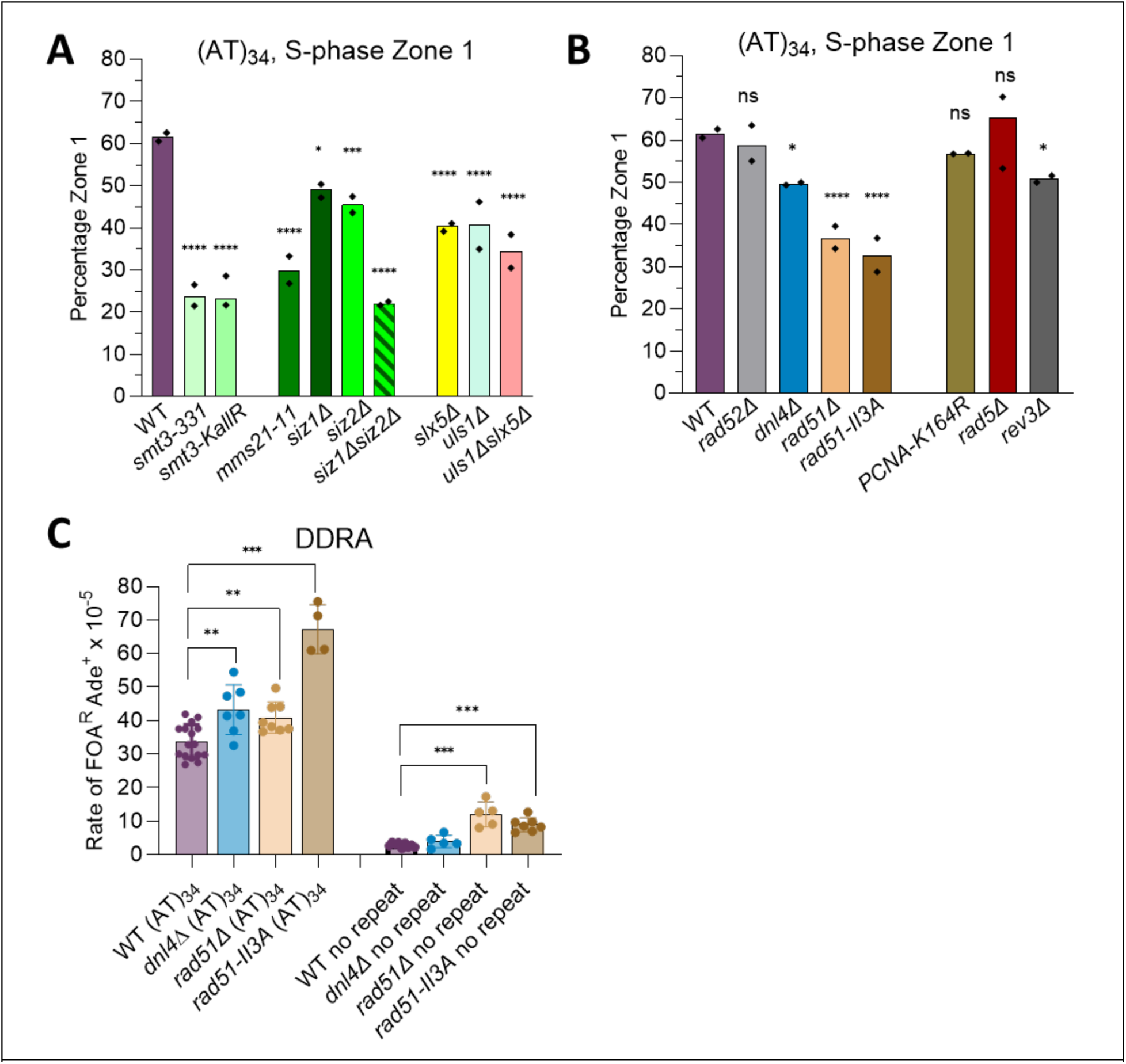
Localization of the (AT)_34_ repeat to the nuclear periphery requires polySUMOylation, STUbL, and recombination activity. A) Percent GFP foci in zone 1 during mid to late S phase for SUMOylation related mutants. B) Percent GFP foci in zone 1 for repair pathway mutants C) Rate of Flex1(AT)_34_-stimulated deletions as measured by the DDRA in repair pathway mutants; points indicate independent experiments. For zoning assays, statistical comparisons by Fisher’s exact test; *p<0.05, ***p<0.001, ****p<0.0001. Zoning data n values are between 90-300 cells per strain; 2 independent isolates were tested for each strain with percents obtained for each indicated by points on bars. Exact n values, zone 1 percentages and p values are listed in Table S3. For DDRAs, pairwise statistical comparison by Welch’s t-test; *p<0.05, **p<0.01, ***p<0.001. Each strain tested with 2 independent isolates. DDRA values and p values are listed in Table S5.

There are three mitotic SUMO E3 ligases in yeast, Mms21/Nse2, Siz1 and Siz2. In the absence of Siz2, Siz2 targets are monoSUMOylated and lose their polySUMOylation chains suggesting that another SUMO E3 ligase, like Mms21, does the initial monoSUMOylation (Horigome et al. 2016; Chung and Zhao 2015; D’Ambrosio and Lavoie 2014). Each mutant *mm21-11*, *siz1Δ,* and *siz2Δ* showed a decrease in localization with the *mm21-11* strain showing the largest decrease (Figure 3A), implicating monoSUMOylation by Mms21 as a key requirement. With expanded (CAG)_130_ repeats mono-SUMOylation is sufficient for relocalization to the NPC. In contrast, there was a dependence on both Mms21 and either Siz1 or Siz2 for peripheral sequestration of a persistent DSB, suggested to be due to sequential action of the ligases with Mms21 mono-SUMOylation occurring first, followed by Siz1/2 polySUMOylation (Horigome et al. 2016). Although Siz1 and Siz2 have different primary targets, they can compensate for each other over time (Cremona et al. 2012). A double *siz1Δsiz2Δ* mutant showed a significant decrease in zone 1 enrichment, to the same level as the *smt3-KallR* mutant (Figure 3A). Therefore, the SUMO ligase requirements for the (AT)_34_ repeat relocalization pathway are more similar to those of an induced DSB than an expanded (CAG)_n_ repeat.

The Slx5/8 SUMO-targeted ubiquitin ligase (STUbL) complex is important for anchoring damaged loci to the NPC, through Slx5 interaction with Nup84. The SUMO interacting motifs (SIMs) on Slx5 were required for anchoring of the (CAG)_130_ collapsed forks to the NPC and reducing their mobility (Su et al. 2015; Whalen et al. 2020). A *slx5Δ* mutant showed a significant decrease in zone 1 localization of the (AT)_34_ repeat compared to the WT to 40% (Figure 3A), suggesting that a SUMO-dependent mechanism is required for the anchoring of the AT repeat to the NPC. We also tested Uls1, another STUbL with SIMs, that inhibits nonhomologous end joining at telomeres and modifies Nup60 of the NPC (Lescasse et al., 2013; Niño et al., 2016). A *uls1Δ* mutant showed a decrease in zone 1 occupancy to 41% while the double *slx5Δuls1Δ* showed a decrease to 34% (Figure 3A). Uls1 was also shown to play a role in peripheral nuclear positioning at a break containing a telomere repeat on one side that inhibits resection (Marcomini et al., 2018). This suggests a role for Uls1 in mediating interaction of poorly resected DSB ends with the nuclear periphery. Interestingly the *slx5Δuls1Δ* double mutant has a viability defect in the presence of DNA damaging agents (Kramarz et al. 2014). Our data indicates that both Slx5 and Uls1 STUbLs play an important role in (AT)_34_ repeat repositioning.

In summary, we conclude that the pathway for (AT)_34_ repeat repositioning to the NPC relies on both mono and polySUMOylation and that both Slx5/8 and Uls1 STUbLs are required.

### Rad51 recombination activity is necessary for the repositioning of the (AT)_34_ repeat to the nuclear periphery

To investigate what repair pathways are required for repositioning to the NPC, mutants in key pathways were tested. The requirement of Rad51 for relocalization of replication barriers to the NP depends on the system. For a (CAG)_130_ repeat, Rad51 is not required for repositioning to the NPC (Whalen et al. 2020), however, for a protein replication fork barrier (RFB) in *S. pombe,* Rad51 and its strand exchange activity are required (Kramarz et al. 2020). For the Flex1(AT)_34_ repeat, a significant decrease in zone 1 repositioning was observed in a *rad51Δ* mutant (Figure 3B). The Rad51 recombinase carries out at least two functions: the formation of the nucleoprotein filament on ssDNA, and the formation of joint molecules between homologous sequences which is essential for strand exchange and D-loop formation (Cloud et al. 2012). To elucidate which of these two functions are required for the relocalization of the AT repeat, a separation of function mutant, *rad51-II3A* was employed. In this mutant the first DNA binding site of Rad51 is intact, allowing it to form a nucleoprotein filament, but the second DNA binding site has three residues mutated to alanine causing it to be defective in strand exchange (Cloud et al. 2012). The *rad51-II3A* mutant exhibited a similar decrease in (AT)_34_ zone 1 percentage as is observed in the absence of Rad51 (Figure 3B), implicating the strand exchange function of Rad51 as necessary for repositioning of the Flex1(AT)_34_-stalled fork to the nuclear periphery.

A deletion of Rad52, required for single-strand annealing, showed no decrease in zone 1 occupancy (Figure 3B). Rad52 also mediates association of Rad51 with ssDNA, replacing RPA, and stabilizes the Rad51 filament against Srs2 dissociation (Ma et al. 2018). Since Rad52 function is not needed for relocation, it may be that RPA or Srs2 binding are inhibited in other ways at this step of the process, allowing enough Rad51 binding to mediate strand exchange. Deletion of Dnl4, the ligase required for both non-homologous and microhomology mediated end joining in *S. cerevisiae*, led to a small but significant decrease in zone 1 repositioning (Figure 3B), suggesting a minor role for end joining in promoting the need for relocation to the NPC.

To test if the post replication repair (PRR) pathway is involved in peripheral positioning of the AT repeat to the nuclear periphery a zoning assay was carried out with a *pol30^K164R^* mutant, which prevents mono and poly-ubiquitination of the K164 residue which is a requirement for translesion systhesis (TLS) and template switch (TS) pathways (Lee and Myung 2008). No defect in zone 1 occupancy was observed in the *pol30^K164R^*mutant (Figure 3B). A deletion of Rad5, required for polyubiquitylation of PCNA at the K164R residue, also had no effect on zone 1 occupancy. This data argues against a requirement for PRR for relocation to the NPC. Interestingly, a deletion of Rev3, the catalytic subunit of Pol ζ, showed a small but significant decrease in zone 1 occupancy (Figure 3B). Since we previously showed that Pol ζ was important for replicating the Flex1(AT)_34_ repeat, this could indicate a role (though not an absolute requirement) for Pol ζ-mediated replication in order to create the substrate that initiates relocation, for example the substrate that is targeted by the SMR complex.

To evaluate the effect of different repair pathways in preventing and healing breaks at the Flex1(AT)_34_ repeat, DDRAs were employed. In the DDRA, a *dnl4Δ* mutant showed no difference in deletions for the no repeat control construct and a slight increase for the AT repeat (Figure 3C), suggesting that end joining is playing a role in repair of AT repeat-induced breaks. The *rad51Δ* mutant had an increase in deletions for the no repeat control (4.2x), which is the expected phenotype if breaks that are normally healed by HR must now be shunted into the more deleterious SSA pathway (Figure 3C). Surprisingly, the increase in SSA-mediated deletions, though still significant, was more muted for the repeat-containing construct (1.2x) (Figure 3C), suggesting that the repeat may be inhibiting Rad51-mediated HR, perhaps due to the predicted hairpin-capped ends. The *rad51II-3A* mutant also exhibited a significant 2-3x increase in SSA events compared to the WT (Figure 3C), supporting the importance of Rad51-dependent strand invasion in preventing (AT)_34_ stimulated deletions. Overall, the analysis of repair pathways revealed a requirement for Rad51-mediated strand exchange in repositioning the AT repeat to the NPC and in protecting it from SSA-mediated deletions.

### Rad51 associates with the Flex1(AT)_34_ repeat in the nuclear interior

To further study the role of Rad51 at the AT repeat and to capture more structural information on the substrate that is repositioned, Rad51 was functionally tagged with the yeast codon optimized yellow fluorescent protein mVenus (Sheff and Thorn 2004), where the mVenus sequence is inserted into a non-conserved internal (i) region of the Rad51 protein (Liu, Mine-Hattab, et al. 2023). In addition to Rad51-i-mVenus, the NP was labeled with mCherry-Nup49 and the LacO array 6.4 kb from the repeat locus with LacI-CFP so that all 3 elements could be visualized in the same cell (Figure 4B). To verify that Rad51-i-mVenus is functioning like the untagged protein, a zoning assay was carried out, and no difference in (AT)_34_ locus positioning compared to WT cells was observed (Figure 4A). In previous studies, spontaneous Rad51 foci were observed in about 5% of cells, either using a functionally tagged Rad51-i-GFP construct (Liu, Mine-Hattab, et al. 2023) or a nonfunctional C-terminally tagged Rad51 in the presence of WT non-tagged Rad51 (Waterman et al. 2019). We scored Rad51-i-mVenus foci in an asynchronous culture and identified mid to late S phase cells by bud and nucleus morphology. The percentage of cells with a spontaneous Rad51 focus is ∼7% for cells containing the no-repeat control sequence (Figure 4C). Interestingly, in WT cells containing Flex1(AT)_34_ the percent of cells with spontaneous Rad51 foci was significantly higher at roughly ∼15%, with the majority of cells exhibiting a single focus (Figure 4C; Table S6). After induction of a DSB with an I-SceI endonuclease, some functionally tagged Rad51 filaments were organized into long rods (Liu, Mine-Hattab, et al. 2023). In contrast, very few Rad51 rods were detected among the spontaneous Rad51 foci (Figure 4C; Table S6). This difference could be due to the different characteristics of a DSB derived from Mus81-Mms4 cleavage of a structure forming AT repeat compared to an HO endonuclease induced break which undergoes extensive resection.

**Figure 4:**
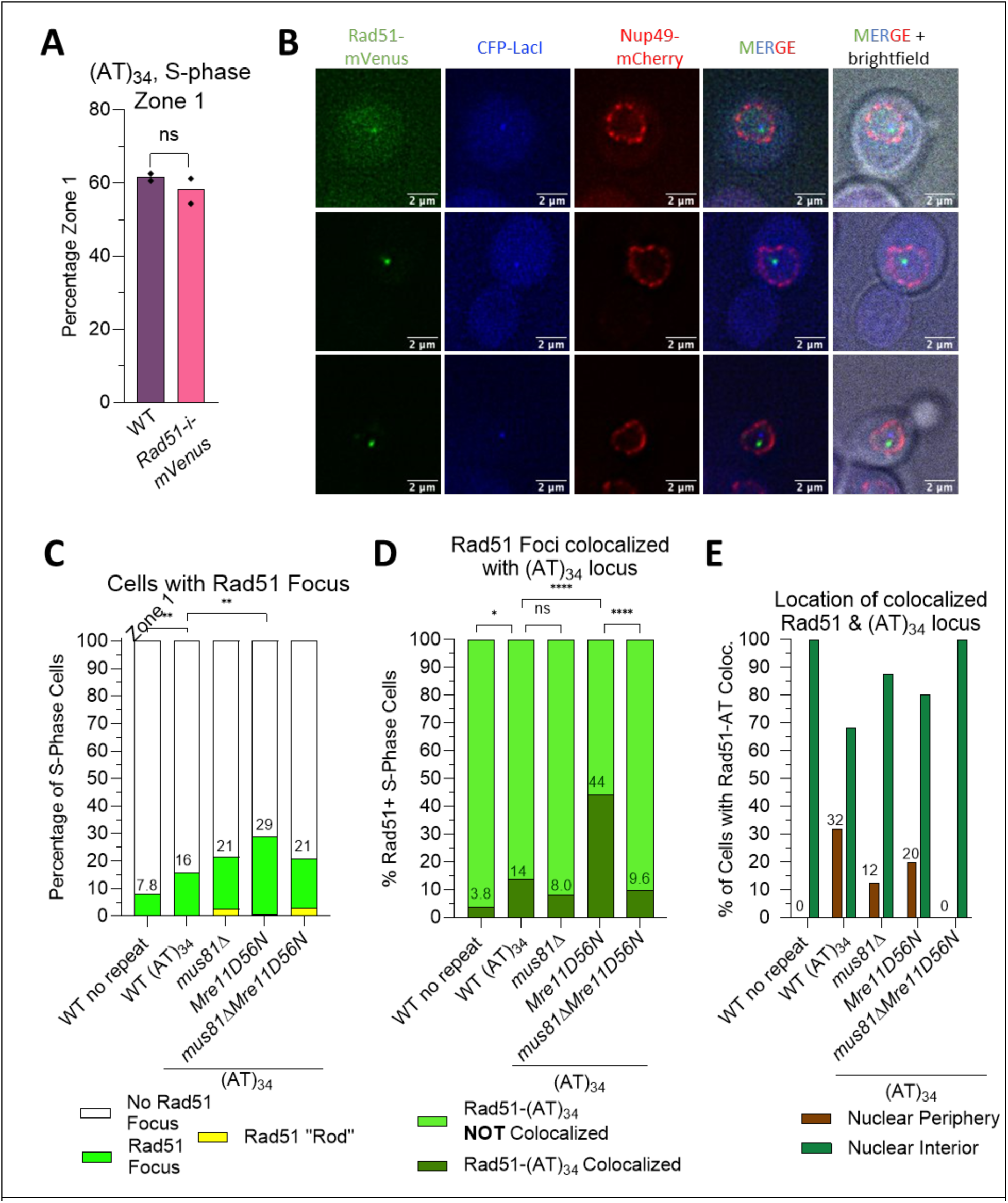
Rad51 colocalizes with the Flex1(AT)_34_ repeat prior to relocalization, dependent on Mus81 cleavage. A) The percentage GFP/CFP foci in zone 1 during mid to late S phase for a strain containing (AT)_34_ on ChVI in a WT strain or a strain with a Rad51 protein containing an internal mVenus tag (Rad51-i-mVenus). B) Sample images of a Rad51-i-mVenus focus colocalized (rows 1, 2) and not colocalized (row 3) with a LacO/LacI-CFP focus. mCherry-Nup49 labels the nuclear periphery. C) The percentage of cells in mid to late S phase containing a Rad51 focus in different mutant backgrounds. D) Of cells that have Rad51 foci, the percentage in which Rad51 and the (AT)_34_ or no repeat control locus (marked by LacO/LacI-CFP) are colocalized (within a 0.3 μm distance; see also Figure S3A-E). E) The percentage of colocalization events that occur at the nuclear interior vs the nuclear periphery in various mutant backgrounds. Comparisons by Fisher’s exact test were as follows. For zoning assays, pairwise statistical comparisons by Fisher’s exact test. Zoning data n values are between 190-250 cells per strain and 2 independent isolates were tested for each strain. Exact n values, zone 1 percentages and p values are listed in Table S3. For percentage of cells with Rad51 foci pairwise statistical comparisons by Fisher’s exact test; **p<0.01. Between 150-410 cells were counted per strain. Exact values and p values in Table S6. For colocalization assay statistical comparisons by Fisher’s exact test; *p<0.05, ****p<0.0001. Between 50-200 Rad51 foci counted per strain. Exact amount of Rad51 foci counted, percentages of colocalization, and p values are listed in Table S7 and Table S8.

To see how often Rad51 foci are at the Flex1(AT)_34_ repeat, the number of Rad51 foci that were within 0.3 microns of the lacI-CFP focus were scored (Figure S3A-E). As expected, few of the spontaneous Rad51 foci co-localized with the control sequence (3.8%), but this was increased to 15% for strains with Flex1(AT)_34_ (Figure 4D; Table S7). This number is an estimate since the array is 6.4 kb from the AT repeat but suggests that at least some of the spontaneous Rad51 foci are forming at the AT repeat tract. The Rad51 foci that colocalized with the (AT)_34_ repeat locus were further evaluated for location within the nucleus (Figure 4E and Table S8). The majority of the time, 68%, Rad51 co-localized with the Flex1(AT)_34_ locus in the nuclear interior, consistent with the requirement for Rad51 for relocalization to the NPC. This verifies that Rad51 interacts with the repeat prior to repositioning to the NPC. However, 30% of the time they were co-localized at the NP whereas this was not observed with the no repeat control, suggesting that Rad51 could travel with the cleaved AT tract out to the NPC.

### Rad51 foci formation reveals a role for Mre11 in processing of AT/TA repeat structures

To understand the order of the processing events, we investigated the requirement for the nucleases that act at the AT repeat for Rad51 focus formation. Rad51 colocalization with the repeat drops from 15% in WT to 8% in *mus81Δ* cells (Figure 4D; Table S7). Though this difference did not reach statistical significance, it was not observed for spontaneous foci not associated with the AT repeat and suggested that Mus81 cleavage might be necessary to create the structure that Rad51 colocalizes with. Strikingly, the nuclease dead *Mre11D56N* mutant showed a significant increase of Rad51 foci co-localizing with the (AT)_34_ locus, to almost 45% of Rad51 foci (Figure 4D; Table S7). This data suggests that in the absence of Mre11 nuclease activity there is a persistence of Rad51 at the repeat tract, consistent with Mre11-dependent processing being required for resolution of a structure. Remarkably, the increase in Rad51 colocalization with the (AT)_34_ locus is dependent on Mus81 as it drops down to 9.6% in the double *mus81ΔMre11D56N* mutant, which is roughly equivalent to the *mus81Δ* single mutant level (Figure 4D; Table S7). These data are most consistent with Mus81 acting before Mre11 to cleave the (AT)_34_ repeat, followed by Rad51 association, and then Mre11 nuclease-dependent resolution of the break.

### The (AT)_34_ repeat structure may disfavor RPA binding

RPA has a high affinity for ssDNA and is involved in stalled fork, gap, and DSB repair pathways (Fanning, Klimovich, and Nager 2006). To gain insight into when and where ssDNA is forming at the AT/TA repeat tract, the C-terminus of Rfa3, the small subunit of yeast RPA, was tagged with mVenus, allowing for live cell colocalization assays of RPA and the (AT)_34_ locus. In previous studies, Rfa3 foci were seen in about ∼20% of untreated cells (Ramonatxo and Moriel-Carretero 2021). Surprisingly, similar rates of Rfa3 foci are seen in both the no repeat control and the (AT)_34_ strains at 22.4% and 18.6% respectively (Figure S4A). The amount of Rfa3 foci that are colocalized with the (AT)_34_ repeat locus (within 0.3 microns of the LacI-CFP focus) was 6% which was not significantly different than the no repeat control locus value of 4.4% (Figure S4B; Table S10). When the RPA and repeat locus foci are colocalized, this occurs more frequently at the nuclear interior (93.3%) than at the nuclear periphery (6.7%) (Figure S4C). The lack of Rfa3 foci associated with the AT repeat locus compared to what was observed for Rad51 foci despite more breaks occurring there, suggests that RPA may be obstructed from binding at the repeat, perhaps due to DNA structure formation.

### The Flex1(AT)_34_ repeat does not result in a converging fork rescue or a Rad52-dependent restart pathway

To further understand replication fork dynamics at the Flex1(AT)_n_ repeat, Okazaki fragment sequencing (Ok-seq) was performed. In the Ok-seq method DNA ligase 1 is conditionally depleted resulting in the accumulation of unligated Okazaki fragments, which are enriched and sequenced to determine the predominant direction of replication (Figure S2A) (Smith and Whitehouse 2012; Smith, Yadav, and Whitehouse 2015). Appropriate strains were created and Ok-seq profiles were compared for the non-structure forming no repeat control sequence and the structure forming Flex1(AT)_34_ repeat sequence, which are inserted ∼6 kb from ARS607 (Figure 1A). ARS607 is a highly efficient early firing origin while ARS608 is an extremely inefficient origin (Shirahige et al. 1993). We observed that the inserted sequences were predominantly replicated by replication forks originating from the ARS607 origin as most sequencing reads align to the Watson strand (Figure 5A; Figure S5A). No differences were detected between ARS608 usage or replication fork direction with either inserted sequence, and there is no evidence of an increase in dormant origin usage or increased converging replication forks due to the (AT)_34_ repeat (Figure 5A; Figure S5B, S5C). We were also not able to detect an (AT)_34_-specific fork stall, which would appear as an accumulation of reads around the AT repeat (Figure 5A; Figure S5B). Fork stalling might not be detectable either due to it being too transient or not occurring in enough cells or in a synchronous-enough manner to detect by this method.

**Figure 5:**
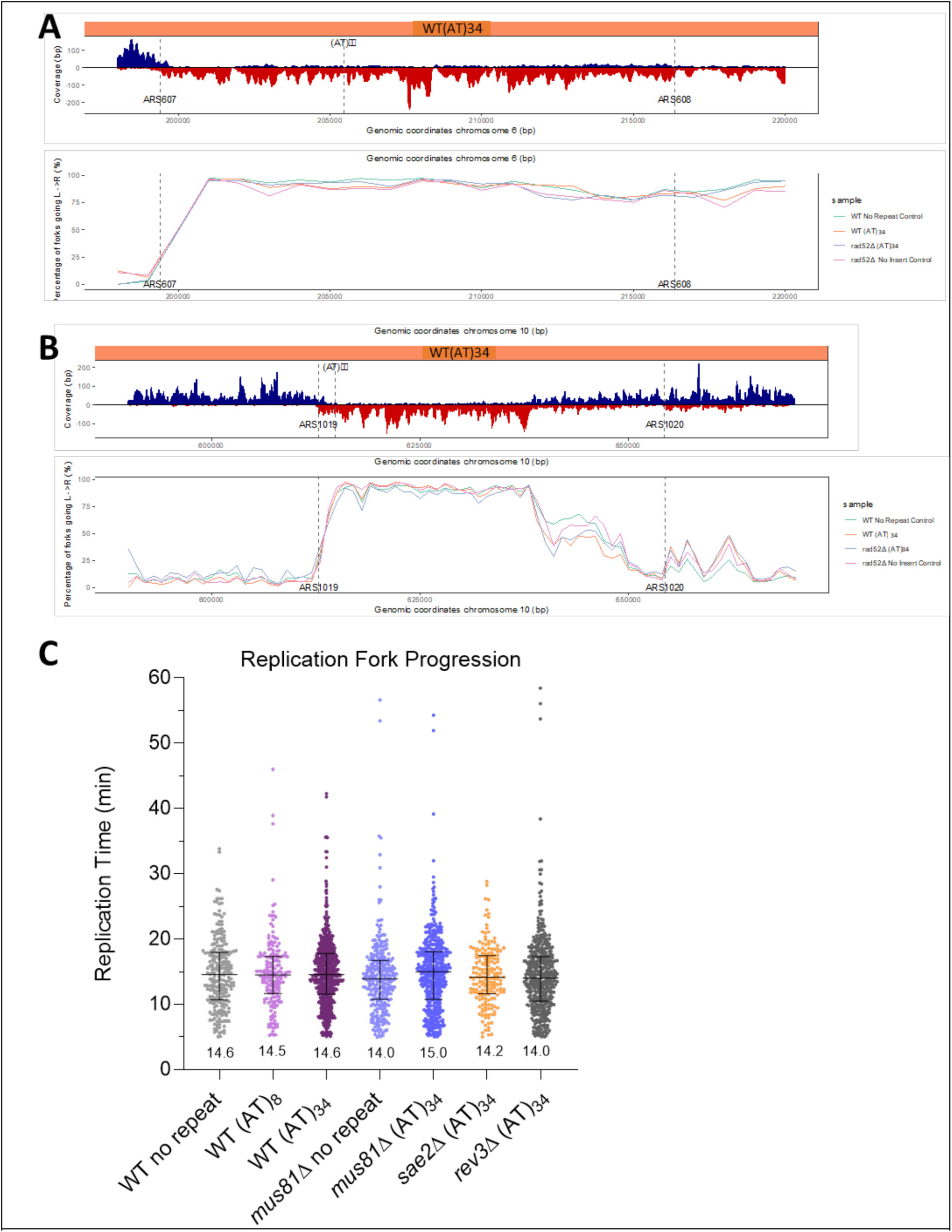
Replication profile of the Flex1(AT)_34_ locus by Okazaki fragment sequencing and replication fork progression by live-cell imaging. A) Okazaki fragment sequencing data for strains with the no repeat control sequence (an AT rich sequence from FRA16D) or the Flex1(AT)_34_ sequence inserted 6 kb to the right of ARS607 in the WT background, or Flex1(AT)_34_ or no insert in the *rad52Δ* mutant background. The percentage of replication forks moving left to right from ARS607 is plotted for the indicated strains. A sample trace of sequencing coverage for the Watson and Crick strands is shown for the WT Flex1(AT)_34_ strain; traces for other strains shown in Figure S5. B) Okazaki fragment sequencing data for either WT or *rad52*Δ strains with either no insert or Flex1(AT)_34_ inserted 7 kb to the right of ARS1019. The percentage of replication forks moving left to right from ARS1019 is shown, along with an example trace for the WT Flex1(AT)_34_ strain. Full chromosome Okazaki fragment sequencing results can be found in Figure S5. C) Replication fork progression through Flex1(AT)n repeats is not delayed in WT or mutant strains. Replication fork progression through Flex1(AT)_8_, Flex1(AT)_34_ or no repeat control are measured for WT and mutant strains. Replication times represent the time required to replicate a ∼22 kb region of Chr IV with or without the Flex1(AT) repeats (Figure S6). Each point represents a single cell measurement and the number underneath the data represents the median value. No statistically significant differences less than p<0.01were observed between the strains. Individual experiments are shown in Figure S6.

The next origin after ARS608 is ARS609, an inefficient origin which is 56.7 kb away from ARS607, which might have impaired our detection of converging forks. To analyze the replication profile in another chromosomal location, we inserted the Flex1(AT)_34_ sequence 7 kb away from ARS1019, a region with more evenly spaced efficient origins. Once again, we did not observe a repeat-dependent change in the replication profile, suggesting that the repeat did not cause a persistently stalled fork at this chromosomal location either (Figure 5B; Figure S5D, S5E).

One way that stalled forks can restart is through a recombination-mediated pathway, which is dependent on Rad52 in *S. pombe* (Lambert et al. 2010). In *S. cerevisiae*, the Rad52 protein is required for all types of recombination, therefore this HR-dependent restart pathway should not be operative in a strain lacking Rad52. When the Ok-seq experiment was done in a *rad52Δ* strain, the replication profile around ARS607 was not markedly altered and no shift in the site of fork convergence between ARS1019 and ARS1020 was observed (Figure 5; Figure S5).

We decided to investigate the replication profile of the Flex1(AT)_34_ repeat by a second method that measures replication fork progression *in vivo* in single cells. This live-cell imaging approach utilizes the lacO and tetO fluorescently labeled arrays that are ∼22 kb apart and normally replicated by *ARS413* (Figure S6A). Replication of the arrays increases their dot fluorescence intensity due to the recruitment of additional fluorescently labeled repressors. By measuring the time difference between the lacO and tetO array duplication events replication fork progression from *ARS413* can be quantitatively measured (Dovrat et al. 2018). For the examination of replication fork progression through the Flex1(AT) repeat tracts, we integrated these repeats between the lacO and tetO arrays followed by live-cell microscopy experiments. Previously, we have used a similar strategy to detect replication slowdown through G4 sequences (Dahan et al. 2018) or GAA repeats (Masnovo et al. 2024). Surprisingly, we observed no statistically significant differences in replication progression between strains containing the structure or non-structure forming inserts, indicating the absence of replication slowdown due to the Flex1(AT)_34_ repeat tract. Similarly, deletion of key genes that impacted relocation to the NPC, *mus81Δ, sae2Δ,* or *rev3Δ*, did not lead to replication slowdown due to the presence of the (AT)_34_ insert (Figure 5C; Figure S6B).

Altogether, we conclude that the Flex1(AT)_34_ repeat is not causing a persistent fork stall on the chromosome, at least not one long-lived enough to be detected by either of the methods used, and we found no evidence for Rad52-dependent fork restart or rescue by a converging fork. Since the replication fork is expected to reach the AT repeat near ARS607 in early S phase, yet relocation to the nuclear periphery doesn’t occur until late S phase, a significant time lag, our data indicate that a persistently stalled fork is not likely to be the structure that provokes relocation. In addition, since Okazaki fragment sequencing revealed no evidence of a converging replication fork at the structure-forming AT repeat, we presume that a converging fork is not the structure that provokes relocation to the nuclear periphery. The data suggest the possibility that the AT repeat is bypassed by the replication fork, but that in the process a structure is left behind that initiates relocation to the nuclear periphery.

## Discussion

In this work, we show that a structure-forming AT repeat relocalizes to the NPC in a replication-dependent manner. This repositioning is dependent on polySUMOylation, the action of SSEs, and the strand exchange function of Rad51. Interruption of this relocalization pathway leads to increased genome instability at the AT repeat as determined by (AT)_34_ stimulated deletions. For relocalization to occur, the AT repeat must reach the length in which a cruciform structure can be formed. This aligns with the result that only Flex1(AT)_23_ and Flex1(AT)_34_ cause an SMR-dependent increase in fragility (Kaushal et al. 2019). Chromosome shattering at (AT/TA)_n_ repeats within common fragile sites and palindromic AT-rich repeats was also shown to be dependent on both AT repeat length and MUS81-EME1 in human cell culture (Shastri et al. 2018; van Wietmarschen et al. 2020). Our results add to the body of work that shows that AT repeats become problematic to replicate and repair at lengths in which they can form a cruciform DNA secondary structure and reveal that relocation to the NPC is a mechanism that maintains genomic stability at structure-forming AT repeats.

Mus81-Mms4 and other components of the SMR complex were shown to be needed for the relocalization of the Flex1(AT)_34_ repeat to the NPC. In theory, Mus81-Mms4 could be cleaving a variety of structures at a stalled or reversed fork, but the additional requirement for the Slx1-Slx4 nuclease and our previous data showing a sharp AT length dependence for stimulating fragility as measured by the DDRA and a chromosome end loss assay (Kaushal et al. 2019) are most consistent with cruciform cleavage in a nick and counter-nick mechanism (Figure 6). As Mus81 is not required for the relocalization of the (CAG)_130_ repeat (Figure 4B) the structure cleaved is specific to the AT repeat. Mms4 phosphorylation, which activates the Mus81-Mms4 nuclease, occurs 30-45 minutes after G1 release, persisting through ∼75 minutes (Waizenegger et al. 2020) consistent with the timing of Flex1(AT)_34_ repositioning.

**Figure 6:**
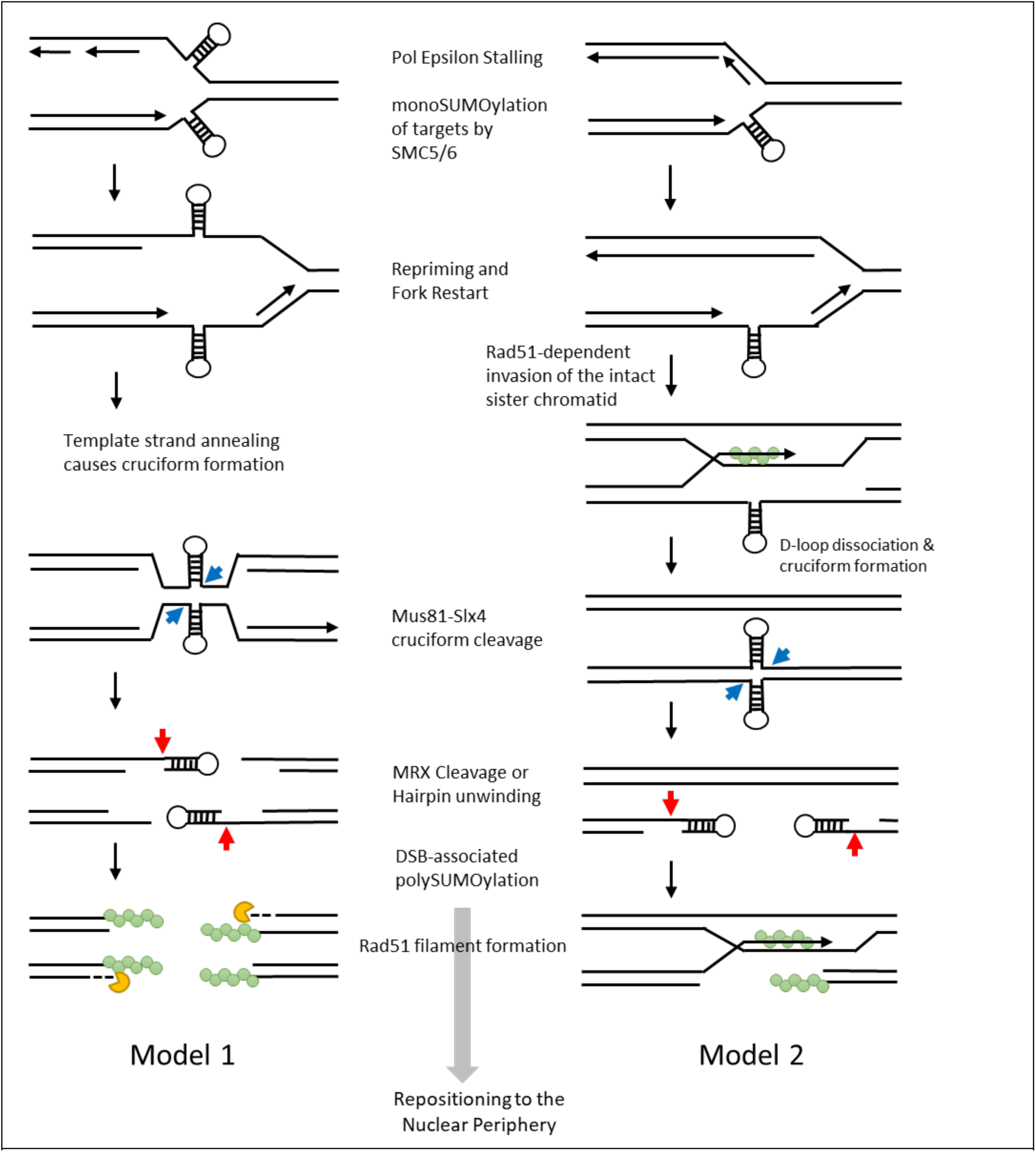
Model for AT/TA structure-mediated fork stalling, cruciform processing, and repositioning to the nuclear periphery. Based on prior data showing Pol epsilon stalling at the AT hairpin base (Das et al. 2024), synthesis of the leading strand is predicted to stall at a hairpin structure. Our replication progression data is consistent with a fork restart event, such as by repriming, occurring before relocation to the NPC, which could result in a gap with a template hairpin. MonoSUMOylation of substrates mediated by Mms21/Nse2 of SMC5/6 may occur in concert with fork stalling/remodeling. In Model 1, a complementary gap on the lagging strand allows strand annealing of the unreplicated regions and cruciform formation. The Mus81-Mms4-Slx4-Slx1 nuclease complex cleaves the cruciform structure, resulting in a break on both sister chromatids, and deletion of the repeat tract after processing of the ends, as is observed in the DDRA. In Model 2, the lagging strand is replicated, providing a template for repair of the hairpin-containing leading strand gap by Rad51-mediated HR (potentially facilitated by Pol ζ). Cruciform formation could occur upon strand dissociation and second end capture, providing the substrate for SLX4 complex SSE cleavage. This would result in a DSB on one chromatid with two hairpin-capped ends. PolySUMOylation of target proteins by Siz1/2 is hypothesized to occur after DSB formation. Hairpin-capped ends could be processed by Sae2/Mre11 or Rad1/Rad10 cleavage or hairpin unwinding coupled to RPA or Rad51 stabilization of the unwound form could occur. In either case, initial Rad51-dependent strand invasion occurs prior to repositioning. Finally, the DSB is repositioned to the NP for repair, mediated by the Slx5/8 or Uls1 STUbL interacting with polySUMOylated proteins and NPC-associated proteins. The completion of repair by copying from the intact template strand, which could occur at the NPC, would result in retention of the AT/TA repeat.

Surprisingly, analysis of fork direction and replication fork progression through both OK-seq and live cell imaging approaches revealed that the repeat does not stall the fork for a length consistent with the timing of the repositioning, which begins about 40-55 minutes after release from G1 arrest and peaks in late S phase, 60-75 minutes after release. These results suggest that replication causes the structure that is targeted by the SLX4 nuclease complex, rather than there being a persistently stalled fork that relocates to the NPC. Using *in vitro* polymerase extension assays, we previously showed that Pol ε is severely stalled by the Flex1(AT)_34_ hairpin structure formed in single-stranded DNA (Das et al. 2024). We also recently determined that the relocation of a (CAG/CTG)_130_ tract is dependent on the phosphorylation of Mrc1, which occurs upon fork uncoupling (Maclay et al. 2025). Together with the lack of an effect of deleting Rad52 on fork progression, our data are consistent with a model in which Pol ε stalls at an AT hairpin, causing fork uncoupling, followed by CMG bypass of the hairpin to allow repriming and fork restart (Figure 6). This would leave behind a ssDNA gap containing the unresolved hairpin on the leading strand. If ssDNA is also present on the lagging strand, the special characteristics of AT repeats could favor annealing to form a cruciform, especially if one or both strands had already formed a hairpin structure (Figure 6, model 1). The cruciform would then provide a substrate for SMR cleavage. Though this model doesn’t require fork reversal, template strand annealing is a potential first step of fork reversal (Liu, Saito, et al. 2023) and annealing of nascent strands could occur behind the cruciform to form another 4-way junction and DSB end. We note that this pathway would lead to deletion of the AT repeat, which we observed when selecting for such events in the DDRA. However, in non-selective conditions the AT repeat is usually maintained. Thus, there must also be other pathways, for example involving cruciform or hairpin unwinding, that can resolve the AT-dependent structure. One possibility is that hairpins formed on the lagging strand are able to be replicated by Pol δ (Das et al. 2024), providing a template for repair of the leading strand gap (Figure 6, model 2). Cruciform formation during this repair event would lead to a structure on only one sister chromatid. This would create two hairpin-capped ends after SMR cleavage, which could preclude RPA binding, as observed, and be difficult to repair, requiring the NPC relocation pathway. Our key observations are (1) a DNA structure that depends on AT length and replication becomes a target of Mus81-Mms4/Slx1-Slx4 cleavage and (2) the repositioning to the nuclear periphery dependent on this cleavage is uncoupled in time from the advancing fork. However, the exact structure targeted and when it occurs in relation to replication remains to be determined. Accounting for background levels of zone 1 occupancy, only about a third of chromosomes containing the AT repeat reposition to the nuclear periphery. Therefore, other pathways to facilitate replication through the DNA structure barrier likely exist, which could include hairpin unwinding, synthesis through the hairpin by Pol ζ, or fork restart mechanisms involving fork reversal or template switching.

A prediction of cleavage at the cruciform base is the production of hairpin-capped ends (Figure 6). Sae2-Mre11 has a hairpin processing function (Lobachev, Gordenin, and Resnick 2002) and could clip off the hairpin-capped ends. The Rad1-Rad10 (human XPF-ERCC1) nuclease has been shown to cleave at the base of hairpin loops (Lu et al. 2015) and thus may also contribute to the hairpin processing. In model 1, the other two ends would not be hairpin capped, and would provide a binding platform for Rad51. We observed an accumulation of Rad51 foci in the *Mre11D56N* nuclease dead mutant that were dependent on Mus81, consistent with a longer or more persistent Rad51 filament, for example on the non-hairpin capped strands, in the absence of Mre11 processing. The zoning and DDRA data both revealed an additive effect of deleting both Mus81 and Sae2, indicating that the Mre11 nuclease may have an additional role independent of Mus81 action at the AT cruciform. The Mre11 nuclease is also known to process reversed fork ends and 3’ ends of nascent strand gaps (step 3 of Model 2) which could lead to ssDNA and provide a Rad51 binding platform (Hale et al. 2023).

After SMR action to create the single chromatid break, hairpin processing or unwinding would be required to create a free 3’ end for strand invasion. The lower-than-expected level of RPA association with the AT repeat may mean that these ends are a poor substrate for RPA binding. This could potentially explain the lack of requirement for Rad52, which normally facilitates RPA-Rad51 exchange, for the repositioning step. The Rad51 protein only needs between 6 and 9 nucleotides to bind (Zaitseva, Zaitsev, and Kowalczykowski 1999) and might be able to coat short 3’ ends made accessible by hairpin unwinding or processing. It is still possible that Rad52 is required for repair at the NPC. A better understanding of NPC-associated repair pathways awaits further characterization.

Mus81-Mms4 and other SMR complex SSEs were shown to contribute to the fragility of *URA3* and *HIS4* inverted repeat palindromes but were not required for the formation of DSBs at *Alu* element inverted repeat palindromes (Ait Saada et al. 2021). The authors suggest that this difference is due to transcription at the palindromic locus. The Flex1(AT)_34_ repeat was inserted in a transcriptionally active region, therefore we cannot rule out a role for transcription in this process. However, there are also size and sequence composition differences between the Flex1(AT)_34_ repeat and these other palindromes which may lead to different abilities to form replication-dependent DNA structures. Long AT repeats were shown to be enriched in regions undergoing mitotic DNA synthesis (MiDAS) by MiDAS-seq, in contrast to Alu elements, long interspersed nuclear elements (LINEs), and short interspersed nuclear elements (SINEs) which were not enriched (Ji et al. 2020). This suggests these other types of palindromic sequences may have different recovery mechanisms.

It was previously shown that monoSUMOylation of targets by the SUMO E3 ligase Mms21 is sufficient for relocalization of an expanded CAG/CTG tract (Whalen et al. 2020). In contrast, the AT repeat requires both Mms21 and Siz1/Siz2 for repositioning, acting more like a persistent double stranded break in its requirements (Horigome et al. 2016). Multiple lines of evidence suggest that the Flex1(AT)_34_ repeat results in a break that is difficult to repair, and one possibility is that polySUMOylation occurs in response to this persistent DSB. PolySUMOylation is also required for the relocalization of a protein mediated replication fork stall in *S. pombe* (Kramarz et al. 2020). Thus, another possibility is that the cruciform-forming (AT)_34_ repeat results in a more persistent fork stall than the hairpin-forming (CAG)_130_ repeat, thus favoring polySUMOylation, but this possibility is less consistent with our Ok-seq and live cell imaging data. A notable observation is that the rise in peripheral association of the (AT)34 repeat occurs over a ∼40 minute time period in S phase (from ∼30-70’ after G1 release) while a (CAG)_130_ tract exhibits a narrower ∼20 minute time window of increasing association (Maclay et al. 2025). Possibly, MonoSUMOylation of RPA and other targets by Mms21 could occur at the initial uncoupling of the replication fork, potentially initiating some early relocation, followed by polySUMOylation by the Siz1 and Siz2 SUMO E3 ligases when a DSB is formed and the ends are resected (Figure 6). The formation of SLX4 condensates in human cells causes compartmentalization of the SUMOylation pathway and promotes SUMOylation of targets including XPF, MUS81, and SLX4 (Alghoul et al. 2023). Therefore, there could be a co-dependent relationship between SUMOylation and nuclease cleavage of structured DNA substrates.

A notable aspect of our model is that it predicts that the AT cruciform will lead to an unreplicated area of the chromosome, which is consistent with what has been observed for CFSs in human cell (Bhowmick et al., 2023). The *yen1Δ* mutant showed a slight increase in the presence of the AT locus at the periphery. It is possible that Yen1 targets AT repeat structures that are left at the periphery at the end of G2. MUS81, SLX4, and GEN1 (the human ortholog of Yen1) were all shown to be needed to visualize gaps at CFSs and loss of any of these nucleases results in reduced MiDAS and reduced ultra-fine anaphase bridges (Benitez et al. 2023; Ying et al. 2013). A GEN1-dependent pathway might be more important in human cells or for larger unreplicated gaps that potentially contain multiple DNA structures or other replication barriers.

MiDAS has been shown to play a role in the maintenance of CFSs such as FRA3B and FRA16D that contain AT rich repeats (Ji et al. 2020). At CFSs, MiDAS requires the SLX4 nuclease scaffold, with MUS81 nuclease activity followed by POLD3 dependent DNA synthesis (Minocherhomji et al. 2015). In addition, MRE11 and CtIP, the mammalian ortholog of Sae2, were found to be essential for MiDAS (Barwacz et al. 2025). As the factors that are required for MiDAS in human cells at CFSs are similar to the requirements of repositioning of the Flex1 (AT)_34_ repeat to the NP, we speculate that an analogous repair pathway, such as break-induced replication (BIR), could be occurring in the yeast nucleus that is facilitated by NPC association. Indeed, studies in yeast have implicated nuclear pore proteins in BIR (Horigome et al. 2016; Chung et al. 2015), and Nup84 was recently shown to play an important role in BIR initiation (Liu et al. 2025). It remains to be determined if NPC association is facilitating BIR at the cleaved AT repeat, but we did find that Pol32, a polymerase subunit essential for BIR, is required to prevent Flex1(AT)_34_ repeat fragility (Das et al. 2024).

Repositioning of persistent DSBs and fork barriers to the nuclear periphery is a conserved mechanism occurring from yeast to *Drosophila* to mammalian cells (Chiolo et al. 2025; Boer et al. 2025). The importance of the NP environment in repair and genome stability is highlighted by the fact that the nuclear periphery can also move towards DSBs (Marnef et al. 2019; Shokrollahi et al. 2024) and DSBs induced in transcriptionally active regions were found to be targeted to the NPC where Rad51 assembly occurs (Le Bozec et al. 2024). It remains to be seen if association with the nuclear periphery plays a role in the repair of CFSs in human cells, and this will be an interesting avenue for future study.

## Data Availability Statement

The data underlying this article are available in the article and in its supplementary material. Raw Okazaki fragment sequencing data is deposited in the SRA database with accession #PRJNA1236355 and processed data will be deposited in the Gene Expression Omnibus (GEO).

## Acknowledgments

We thank Ruben Martinez and Mili Das in the Freudenreich lab for help with strain construction and some DDRAs, Natasha C. Morse for help with initial OK-seq experiments, Sonia Kadyan for help with strain construction, Angela Taddei for sharing the Rad51-i-GFP plasmid, Zaid Salah for constructing the Rad51-i-mVenus plasmid, Xiaolan Zhao for sharing the *smt3-KallR* mutant strain used to introduce the mutant into our strains and Jim Haber for sharing the Rad51-II3A strain used as a template to introduce the mutation into our strains.

## Author Contribution statement

J.L.B., conceptualization, investigation, formal analysis, visualization and writing-original draft. S.M.C., conceptualization, investigation, formal analysis, visualization and writing-review & editing. A.J.K., investigation, formal analysis. D.N.K., investigation, formal analysis. B.T., data curation, formal analysis, visualization. D.D., resources, software. A.A., resources, formal analysis, supervision, funding acquisition. D.J.S., conceptualization, formal analysis, supervision, resources, funding, writing-review & editing. C.H.F., conceptualization, formal analysis, funding acquisition, project administration, supervision, visualization, writing-original draft.

