## Supplemental Tables and Figures for "Cruciform-forming AT/TA repeats are acted upon by structure selective endonucleases and Rad51 prior to repositioning to the nuclear periphery for repair"

**Supplemental Figures**

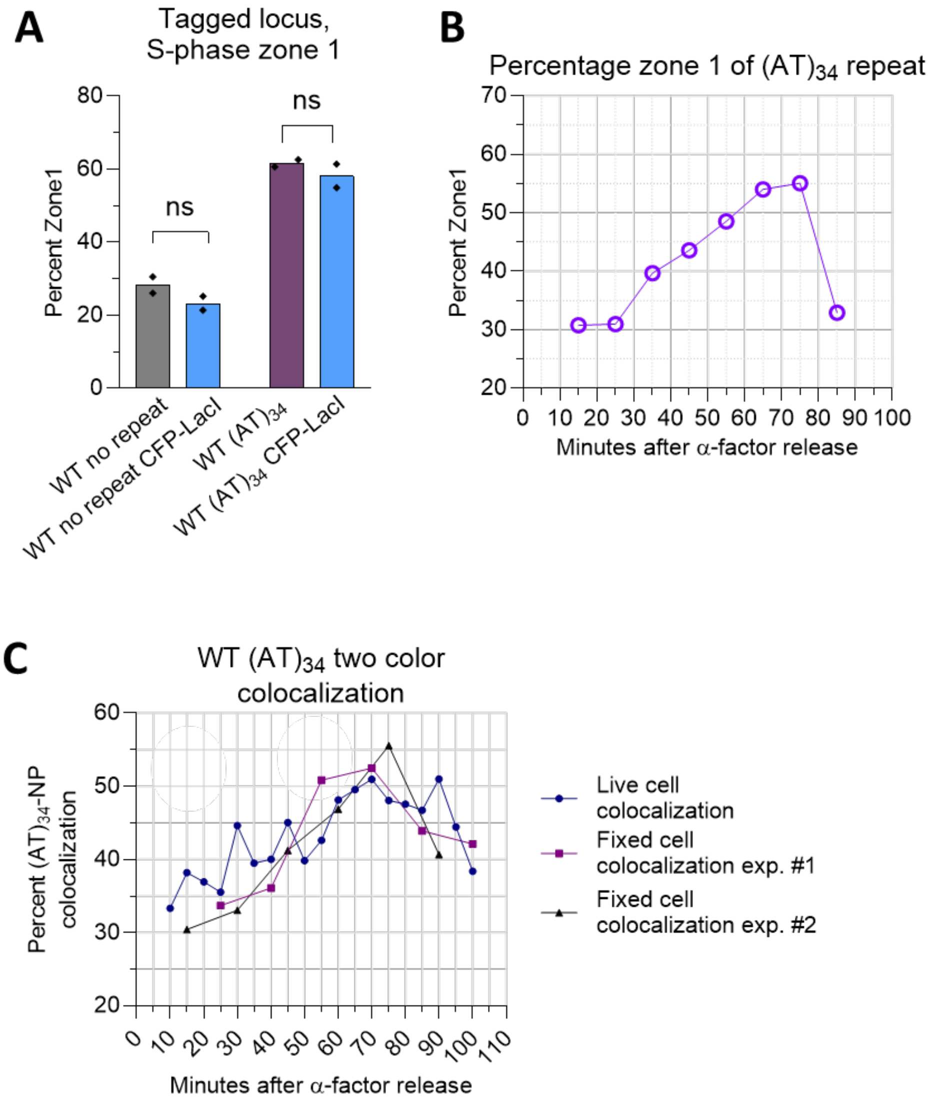

**Figure S1: The structure forming AT/TA repeat relocalizes to the nuclear periphery.** A) The percentage GFP or CFP foci in zone 1 during mid to late S phase for the GFP-LacI/GFP-Nup49 strain and the CFP-LacI/mCherry-Nup49 strain. B) Zoning Analysis of the Flex1$(AT)_{34}$ locus after a-factor arrest in G1 and release into S phase. C) Percentage of colocalization of the $(AT)_{34}$ locus (marked by CFP-LacI/LacO) with the NP after alpha factor arrest in G1 and release into S phase; independent experiments are plotted separately. For 2 experiments, cells taken at each time point were fixed before analysis. For the live cell experiment, cells at each time point were imaged and then analyzed and 3 replicates were combined to reach an n value of ~100 cells per time point. For zoning assays, pairwise statistical comparisons were by Fisher's exact test. Zoning data n values are between 150-210 cells per timepoint; exact n values, zone 1 percentages and p values are listed in Table S3. For colocalization assays n values are between 99 and 189 cells per timepoint; exact n and colocalization values are in Table S4.

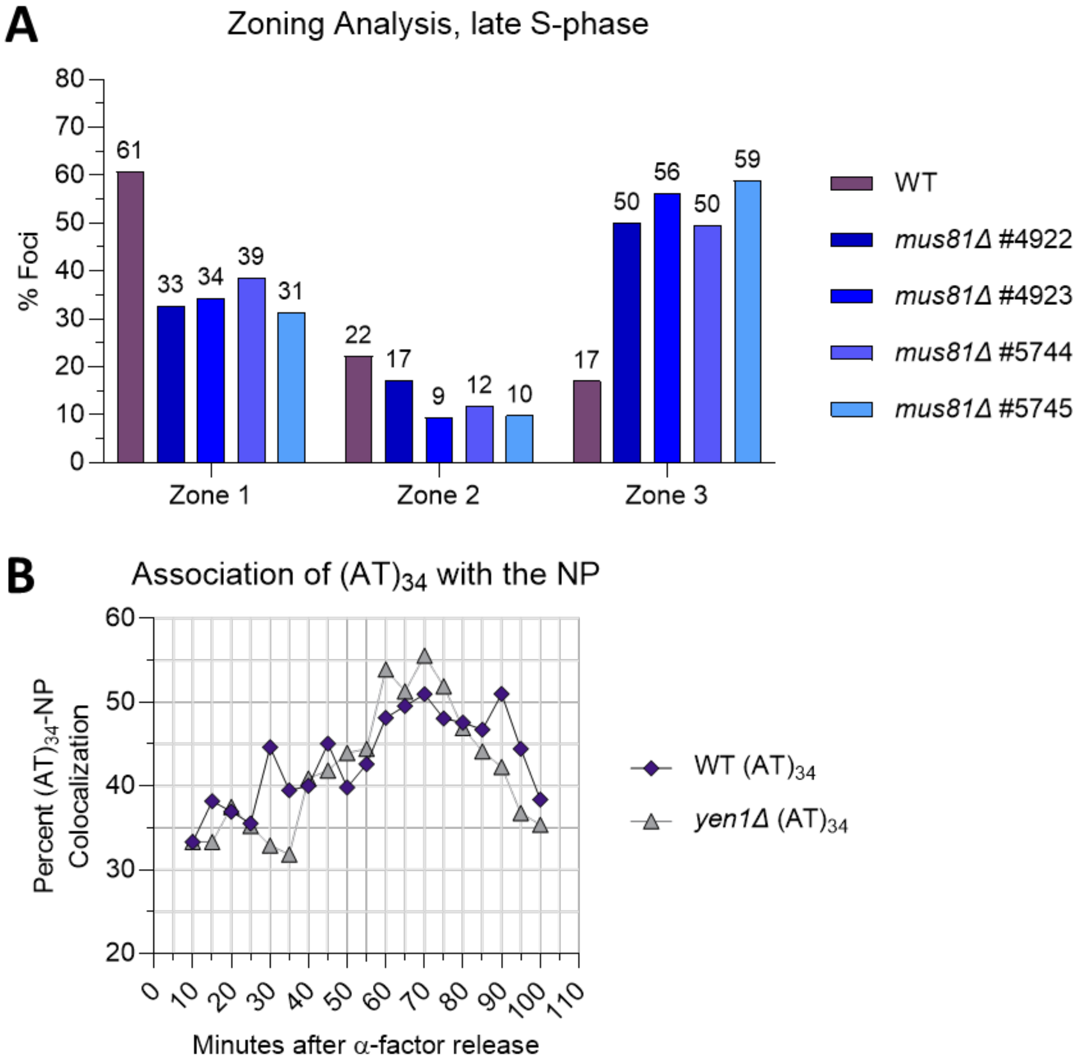

**Figure S2: *Mus81Δ* causes a decrease in zone 1 localization while *yen1Δ* does not have an effect.** A) The percentage GFP-LacI/LacO foci in zone 1, 2 or 3 during mid to late S phase for four independent *mus81Δ* Flex1$(AT)_{34}$ isolates. Zoning data n values are between 30-120 cells per isolate. Exact n values, zone 1 percentages and p values are listed in Table S3. B) Live cell analysis of the percentage of colocalization of the Flex1$(AT)_{34}$ locus with the NP after alpha factor arrest in G1 and release into S phase for WT and *yen1Δ* strains. Cells analyzed for each time point were 99-119 for WT and 63-91 for *yen1Δ*. Exact n and colocalization values seen in Table S4.

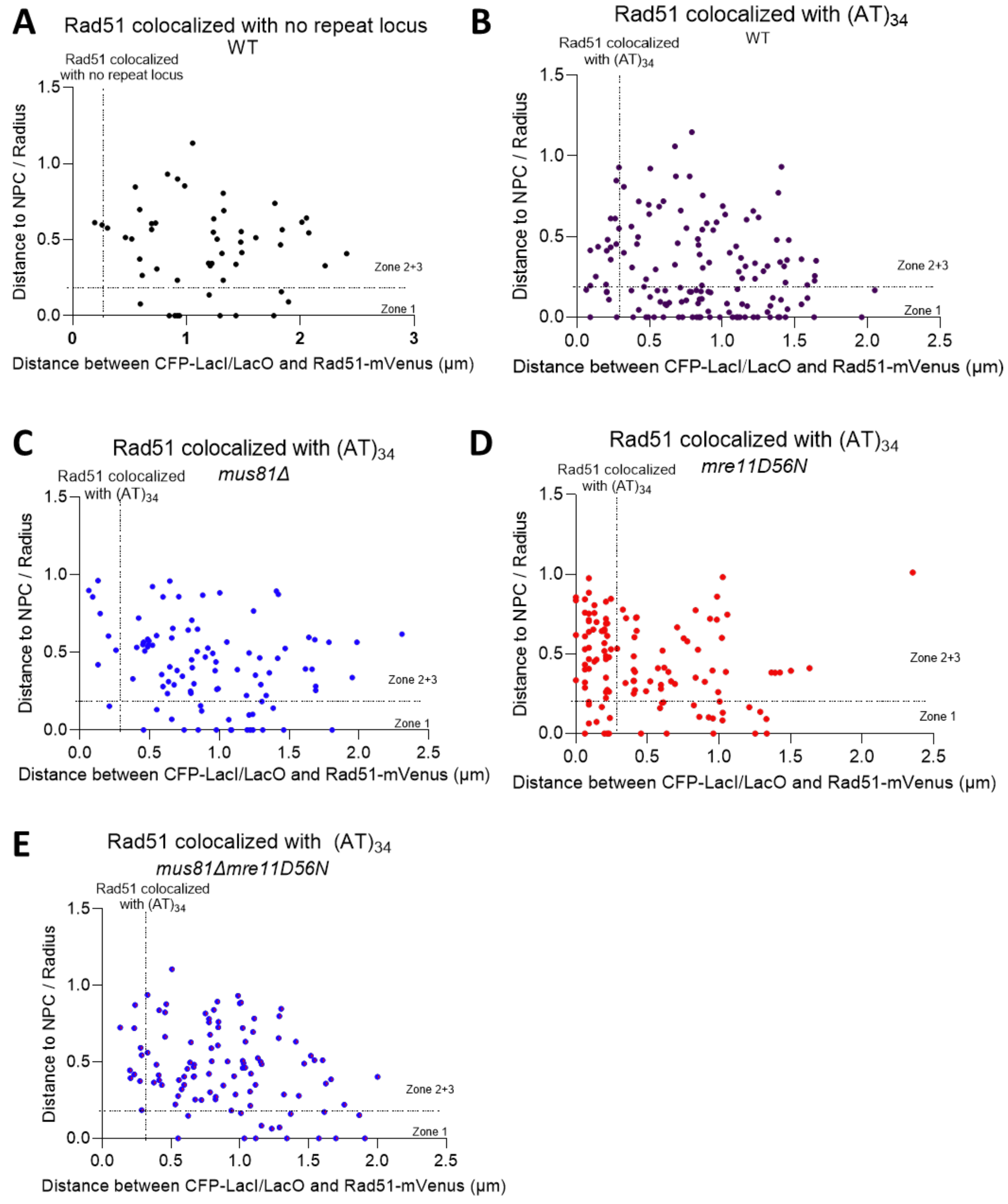

**Figure S3: Distance data for Rad51 colocalization experiments.** A-E) Relative distance of indicated nuclear elements for each mutant. Cutoffs used to score Rad51 foci and the Flex1(AT)$_{34}$ repeat locus (marked by the CFP-LacI/LacO array 6.4 kb away) as co-localized are indicated by the vertical dotted line. Cutoffs used to score the Flex1(AT)$_{34}$ repeat locus as occupying zone 1 by co-localization with the NPC (mCherry-Nup49) are indicated by the horizontal dotted line.

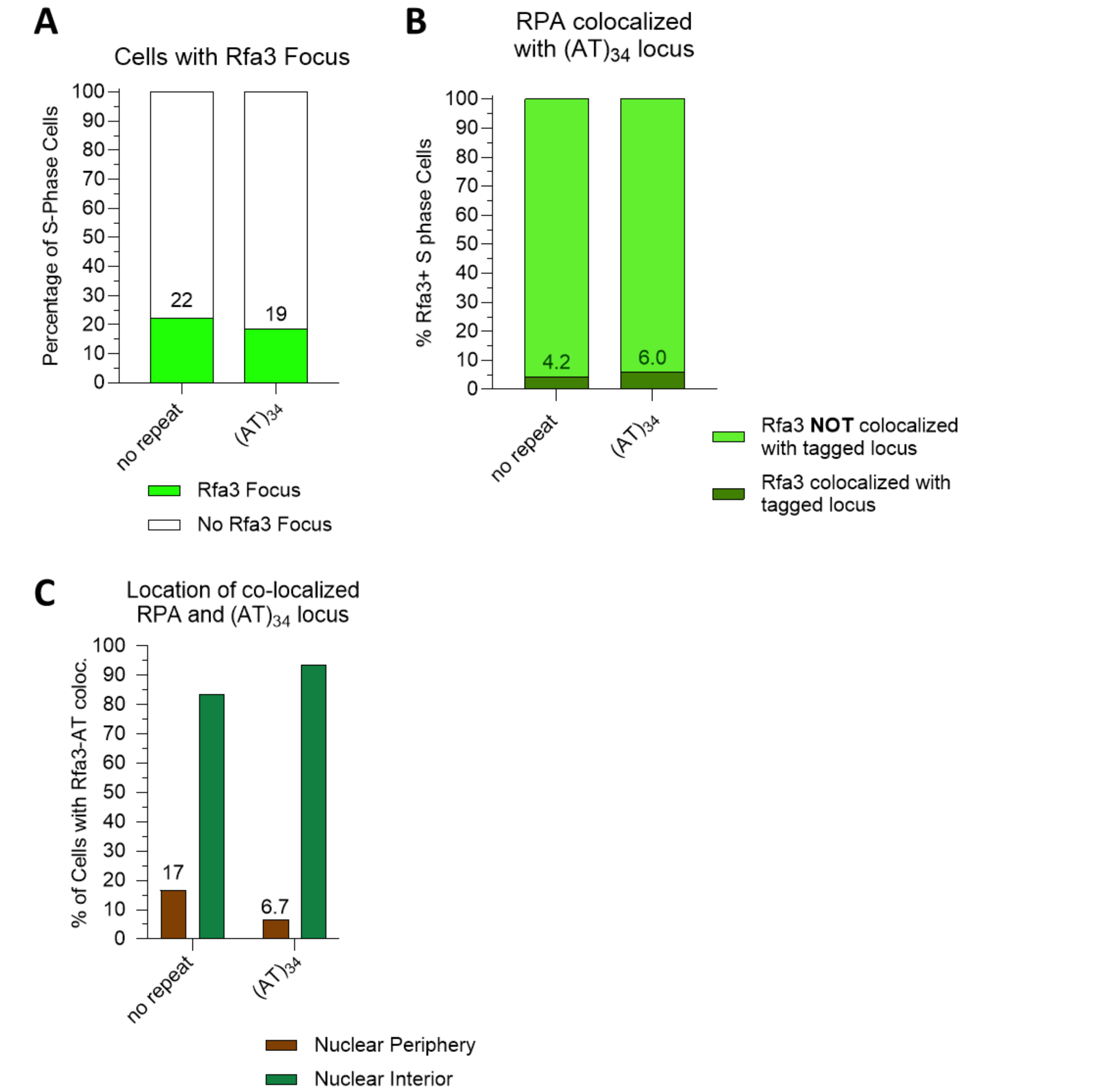

**Figure S4: RPA does not colocalize with Flex1(AT)$_{34}$ repeat more than the no repeat control.** A) The percentage of cells in mid to late S phase containing a Rfa3 focus in different mutant backgrounds. B) Of cells that have Rfa3 foci, the percentage in which Rfa3 and the (AT)$_{34}$ or no repeat locus (marked by LacO/LacI-CFP) are colocalized (within a 0.3 µm distance). C) The percentage of colocalization events that occur at the nuclear interior vs the nuclear periphery. Pairwise comparisons by Fisher's exact test were as follows. For percentage of cells with Rfa3 foci pairwise statistical comparisons by Fisher's exact test. Between 150- 300 cells were counted per condition. Exact values and p values in Table S8. For colocalization assay statistical comparisons by Fisher's exact test. Between 140-250 Rfa3 foci counted per condition. Exact amount of Rfa3 foci counted, percentages of colocalization, and p values are listed in Table S9 and Table S10.

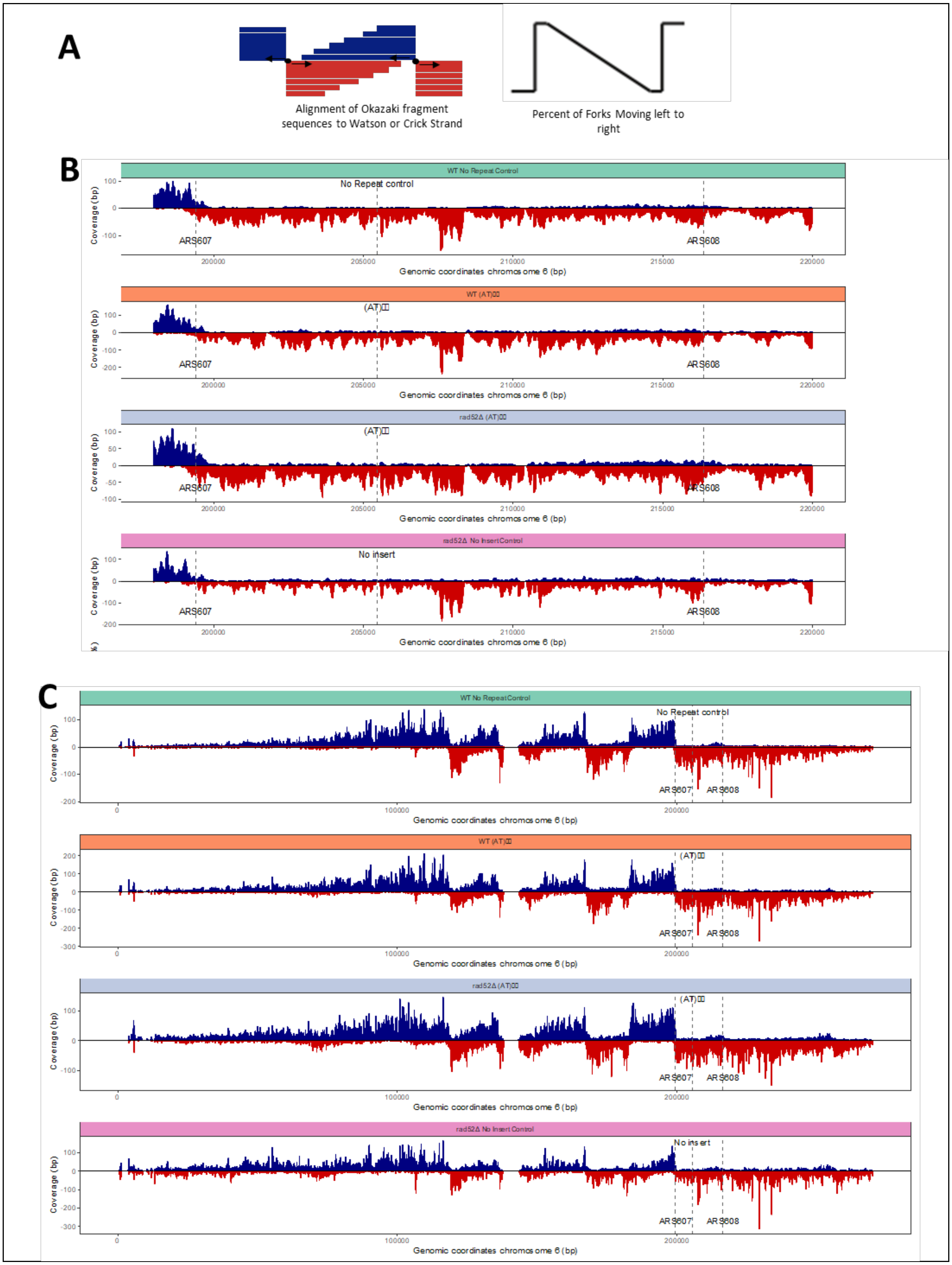

A
Alignment of Okazaki fragment sequences to Watson or Crick Strand
Percent of Forks Moving left to right
B
WT No Repeat Control
No Repeat control
rad52Δ (AT)
rad52Δ No Insert Control
No insert
WT (AT)
ARS607
ARS608
Coverage (bp)
Genomic coordinates chromosome 6 (bp)
C
No Repeat control
No insert

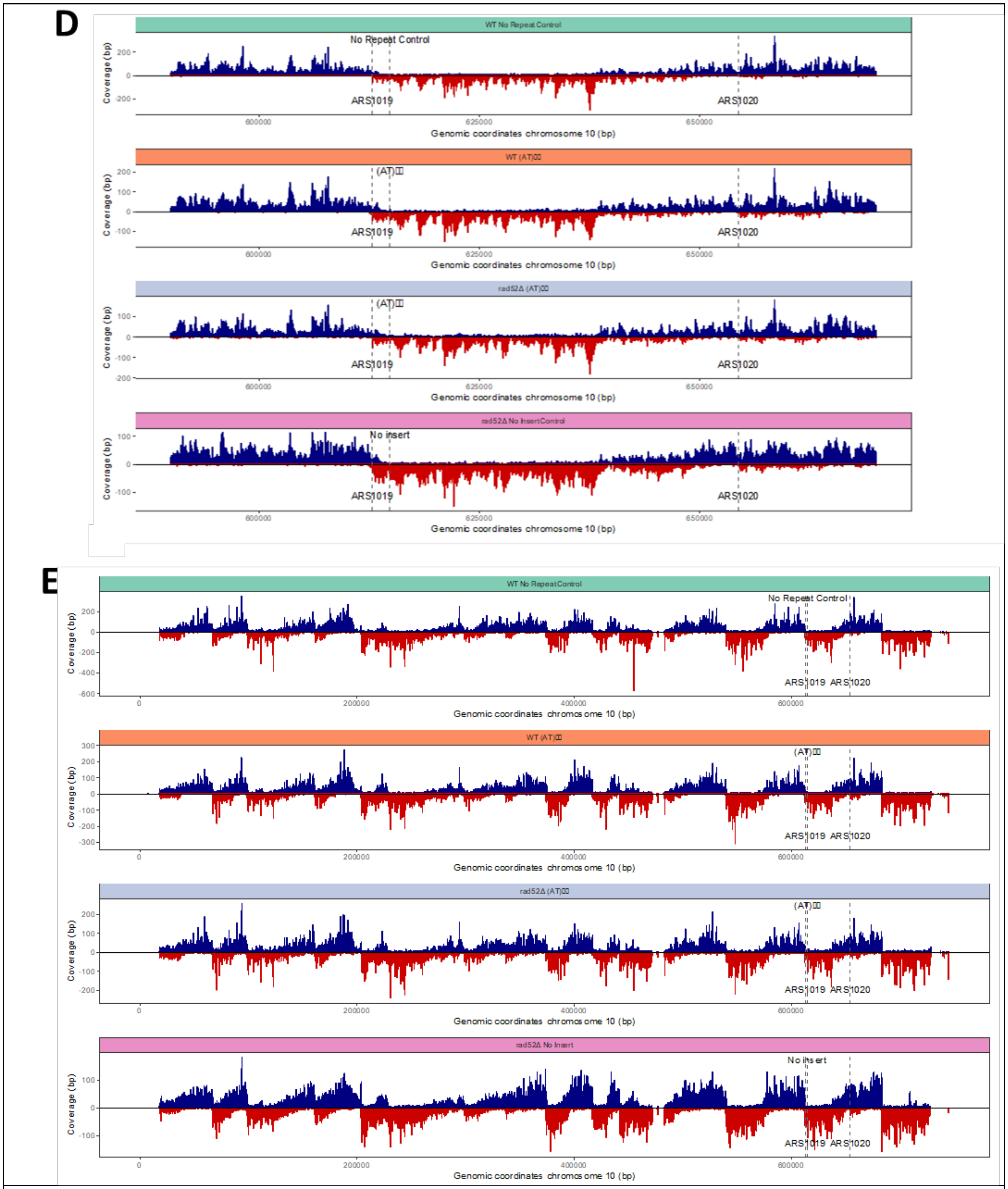

**Figure S5: Okazaki fragment sequencing replication profile of *S. cerevisiae* chromosome 6 and chromosome 10 with indicated inserted elements.** A) Schematic of Okazaki fragment sequencing for data interpretation. Relates to Figure 5. B) Individual Okazaki fragment sequencing results of region of interest for chromosome VI for different inserts and mutant backgrounds. C) Full chromosome Okazaki fragment sequencing results for chromosome VI. The regions of interest shown in Figure 5A and S5B are marked. D) Individual Okazaki fragment sequencing results of the region of interest for chromosome X for different inserts and mutant backgrounds. E) Full chromosome Okazaki fragment sequencing results for chromosome X. The regions of interest shown in Figure 5B and S5D are marked.

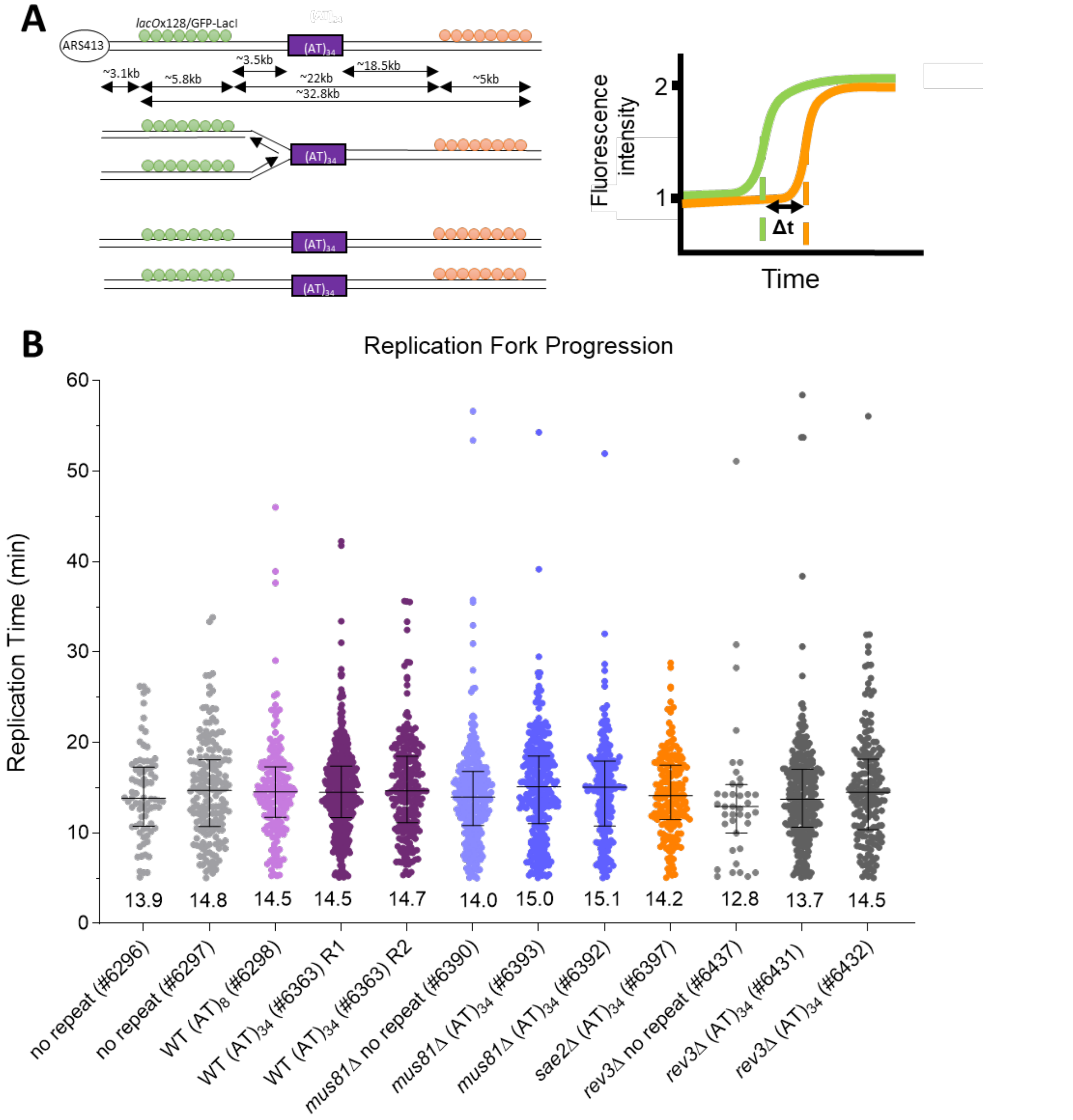

**Figure S6: Individual live-cell imaging replication fork progression assays.** A) Schematic for interpretation of live-cell imaging replication fork progression assay. B) Individual experiments of replication fork progression separated by strain and experiment for the time required to replicate a ~22 kb region of Chr IV with either a no repeat control, Flex1$(AT)_8$ or Flex1$(AT)_{34}$ inserted. Different mutant backgrounds as shown. Each point represents one measured

| |
|---|
| replication event and number underneath the data represents the median values. The bars represent the interquartile range. |

**Supplemental Tables**

**Table S1**: *S. cerevisiae* strains used in this study

| **Strain** | **Description** | **Identifier** |
|---|---|---|
| DDRA assay strains | | |
| *lys2::ADE2* (background: YPH499) | *MATa leu2-Δ1 ura3-52 his3-Δ200 trp1-Δ63 ade2Δ::hisG (salmonella) lys2::ADE2* | CFY #2268<br>(Kaushal et al., 2019) |
| Flex1(AT)$_{34}$ | *lys2::ADE2::URA3-S5'-*Flex1(AT)$_{34}$-S3' (background: CFY #2268) | CFY #2525, #2712<br>(Kaushal et al., 2019) |
| Flex1(AT)$_{34}$ *dnl4Δ* | *dnl4::KANMX*<br>(background: CFY #2525) | CFY #6508 & #6532<br>This study |
| Flex1(AT)$_{34}$ *rad51Δ* | *rad51::NATMX* (background: CFY #2525) | CFY #4705 & #4708<br>This study |
| Flex1(AT)$_{34}$ *rad51-II3A* | *rad51-II3A-KANMX (background: CFY #2525)* | CFY #5783 & #5784<br>This study |
| Flex1(AT)$_{34}$ *rad52Δ* | *rad52::KANMX* (background: CFY #2525) | CFY #5781 & #5782<br>This study |
| Flex1(AT)$_{34}$ *mus81Δ* | *mus81::KANMX* (background: CFY #2525) | CFY #3377 & #3378<br>(Kaushal et al., 2019) |
| Flex1(AT)$_{34}$ sae2*Δ* | *sae2::KANMX* (background: CFY #2525) | CFY #3520 & #3251<br>(Kaushal et al., 2019) |
| Flex1(AT)$_{34}$ *mus81Δsae2Δ* | *sae2::NATMX* (background: CFY #3377) | CFY #5787<br>This study |
| Flex1(AT)$_{34}$ *rad51-II3A* | *rad51-II3A-KANMX*<br>(background: CFY #2525) | CFY #5783 & #5784<br>This study |
| Flex1(AT)$_{34}$ *slx5Δ* | *slx5::KANMX* (background: CFY #2525) | CFY #5711 & #5712<br>This study |
| Flex1(AT)$_{34}$ *nup84Δ* | *nup84::KANMX* (background: CFY #2525) | CFY #5672 & #5673<br>This study |
| Flex1(AT)$_{34}$ *mps3Δ75-150* | *mps3Δ75-150-HPH*<br>(background: CFY #2525) | CFY #5634 & #5635 & #5726 & #5727<br>This study |
| no repeat control | *lys2::ADE2::URA3*-no repeat control (background: CFY #2268) | CFY #2864<br>(Kaushal et al., 2019) |
| no repeat control *dnl4Δ* | *dnl4::KANMX*<br>(background: CFY #2864) | CFY #6510 & #6511<br>This study |
| no repeat control *rad51Δ* | *rad51::NATMX* (background: CFY #2864) | CFY #6077 & #6078<br>This study |

| | | |
|---|---|---|
| no repeat control *rad51-II3A* | *rad51-II3A-KANMX* (background: CFY #2864) | CFY #6542 & #6549 This study |
| Zoning assay strains | | |
| No insert GFP-Nup49 *lacO*/GFP-LacI | *MATa ade2-1 can1-100 his3-11,-15::GFP-LacI:HIS3 trp1-1, ura3-1, leu2-3,-112 nup49::GFP-NUP49 Chr6int1::lacO:4xLexA array:TRP1* (*Chr6int1* is 272 bp upstream of *ARS607,* within *PES4*) | CFY # 2596 (Su et al., 2015) |
| No repeat control GFP-Nup49 *lacO*/GFP-LacI | *Chr6int2::No repeat control-HPH* (Chr6int2 is 6371 bp from Chr6int1 and 560bp upstream of Ta(AGC)F gene) (background: CFY #2596) | CFY #4839 & #4840 This study |
| $(AT)_{14}$ GFP-Nup49 *lacO*/GFP-LacI | *Chr6int2::S5'-Flex1*$(AT)_{14}$*-S3'-HPH* (Chr6int2 is 6371 bp from Chr6int1 and 560bp upstream of Ta(AGC)F gene) (background: CFY #2596) | CFY #4754 & 4755 This study |
| $(AT)_{23}$ GFP-Nup49 *lacO*/GFP-LacI | *Chr6int2::S5'-Flex1*$(AT)_{23}$*-S3'-HPH* (Chr6int2 is 6371 bp from Chr6int1 and 560bp upstream of Ta(AGC)F gene) (background: CFY #2596) | CFY #5067 & #5068 This study |
| $(AT)_{34}$ GFP-Nup49 *lacO*/GFP-LacI | *Chr6int2::S5'-Flex1*$(AT)_{34}$*-S5'-HPH* (Chr6int2 is 6371 bp from Chr6int1 and 560bp upstream of Ta(AGC)F gene) (background: CFY #2596) | CFY #4840 & #4841 This study |
| $(CAG)_{130}$ GFP-Nup49 *lacO*/GFP-LacI | *Chr6int2::(CAG)130* (Chr6int2 is 6371 bp from Chr6int1 and 560bp upstream of Ta(AGC)F gene) (background: CFY #2596) | CFY #4300 (Whalen et al., 2020) |
| $(AT)_{34}$ GFP-Nup49 *lacO*/GFP-LacI (Mat $\alpha$) | Mating #4841x #4300 | CFY #5962 This study |
| *slx5Δ* $(AT)_{34}$ GFP-Nup49 *lacO*/GFP-LacI | *slx5::KANMX* (background: CFY #4841) | CFY #5338 & #5339 This study |
| *uls1Δ* $(AT)_{34}$ GFP-Nup49 *lacO*/GFP-LacI | *uls1::NATMX* (background: CFY #5962) | CFY #6512 & #6513 This study |

| | | |
|---|---|---|
| *uls1Δslx5Δ* $(AT)_{34}$ GFP-Nup49 *lacO*/GFP-LacI | *uls1::NATMX* (background: CFY #5338) | CFY #6536 & #6537<br>This study |
| *smt3-331* $(AT)_{34}$ GFP-Nup49 *lacO*/GFP-LacI | *smt3-331-NATMX* (background: CFY #4841) | CFY #5854 & #5855<br>This study |
| *smt3-KallR* $(AT)_{34}$ GFP-Nup49 *lacO*/GFP-LacI | *smt3-KallR-KANMX* (background: CFY #6157) | CFY #6489 & #6490<br>This study |
| *mms21-11* $(AT)_{34}$ GFP-Nup49 *lacO*/GFP-LacI | *mms21-11-KANMX* (background: CFY #4841) | CFY #5487 & #5488<br>This study |
| *siz1Δ* $(AT)_{34}$ GFP-Nup49 *lacO*/GFP-LacI | *siz1::NATMX* (background: CFY #4841) | CFY #5505 & #5506<br>This study |
| *siz2Δ* $(AT)_{34}$ GFP-Nup49 *lacO*/GFP-LacI | *siz2::KANMX* (background: CFY #4841) | CFY #5507 & #5508<br>This study |
| *siz1Δsiz2Δ* $(AT)_{34}$ GFP-Nup49 *lacO*/GFP-LacI | *siz1::NATMX* (background: CFY #5507) | CFY #5588 & #5589<br>This study |
| *rad52Δ* $(AT)_{34}$ GFP-Nup49 *lacO*/GFP-LacI | *rad52::KANMX* (background: CFY #4841) | CFY #5775 & CFY #5776<br>This study |
| *dnl4Δ* $(AT)_{34}$ GFP-Nup49 *lacO*/GFP-LacI | *dnl4::KANMX* (background: CFY #6184) | CFY #6505 & #6507<br>This study |
| *rad51Δ* $(AT)_{34}$ GFP-Nup49 *lacO*/GFP-LacI | *rad51::KANMX* (background: CFY #4841) | CFY #5445 & #5446<br>This study |
| *rad51-II3A* $(AT)_{34}$ GFP-Nup49 *lacO*/GFP-LacI | *rad51-II3A-KANMX* (background: CFY #4841) | CFY #5785 & #5786<br>This study |
| *PCNA-K164R* $(AT)_{34}$ GFP-Nup49 *lacO*/GFP-LacI | *PCNA-K164R-KANMX* (background: CFY #4841) | CFY #6035 & #6036<br>This study |
| *rad5Δ* $(AT)_{34}$ GFP-Nup49 *lacO*/GFP-LacI | *rad5::KANMX* (background: CFY #4841) | CFY #6005 & #6006<br>This study |
| *rev3Δ* $(AT)_{34}$ GFP-Nup49 *lacO*/GFP-LacI | *rev3::KANMX* (background: CFY #4841) | CFY #6213 & #6248<br>This study |
| *Rfa1-mRuby* $(AT)_{34}$ GFP-Nup49 *lacO*/GFP-LacI | *Rfa1-mRuby-KANMX* (background: CFY #4841) | CFY #5108<br>This study |
| *yen1Δ* $(AT)_{34}$ GFP-Nup49 *lacO*/GFP-LacI | *yen1::NATMX* (background: CFY #5108) | CFY #5150 & #5151<br>This study |
| *slx1Δ* $(AT)_{34}$ GFP-Nup49 *lacO*/GFP-LacI | *slx1::KANMX* (background: CFY #4841) | CFY #6148 & #6149<br>This study |
| *slx4Δ* $(AT)_{34}$ GFP-Nup49 *lacO*/GFP-LacI | *slx4::KANMX* (background: CFY #4841) | CFY #5336, #5931 & #5932<br>This study |
| *mus81Δ* $(AT)_{34}$ GFP-Nup49 *lacO*/GFP-LacI | *mus81::KANMX* (background: CFY #4841) | CFY #4922, #4923, #5744 & #5745<br>This study |
| *mms4Δ* $(AT)_{34}$ GFP-Nup49 *lacO*/GFP-LacI | *mms4::KANMX* (background: CFY #4841) | CFY #5517 & #5518<br>This study |
| *rad1* $(AT)_{34}$ GFP-Nup49 *lacO*/GFP-LacI | *rad1::KANMX* (background: CFY #4841) | CFY #6135 & #6136<br>This study |

| | | |
|---|---|---|
| *Mre11-mRuby2* GFP-Nup49 *lacO*/GFP-LacI | *Mre11-mRuby2-KANMX* (background: CFY #4841) | CFY #5546<br>This study |
| *sae2Δ* $(AT)_{34}$ GFP-Nup49 *lacO*/GFP-LacI | *sae2::NATMX* (background: CFY #5546) | CFY #5597 & #5598<br>This study |
| *mre11-D56N* $(CAG)_{130}$ GFP-Nup49 *lacO*/GFP-LacI | *mre11-D56N, Chr6int2::* $(CAG)_{130}$-*HPH* (Chr6int2 is 6371 bp from Chr6int1 and 560bp upstream of Ta(AGC)F gene) | CFY #4708<br>(Whalen et al., 2020) |
| *mre11D56N* $(AT)_{34}$ GFP-Nup49 *lacO*/GFP-LacI | Mating #4841 x #4708 | CFY #6041, #6042 & #6043<br>This study |
| *mus81Δsae2Δ* $(AT)_{34}$ GFP-Nup49 *lacO*/GFP-LacI | *sae2::NATMX* (background: CFY #5744) | CFY #5796 & #5797<br>This study |
| $(CAG)_{130}$ GFP-Nup49 *lacO*/GFP-LacI | *Chr6int2::*$(CAG)_{130}$*-HPH* (Chr6int2 is 6371 bp from Chr6int1 and 560bp upstream of Ta(AGC)F gene) (background: CFY #2956) | CFY #2744<br>(Su et al., 2015) |
| *mus81Δ* $(CAG)_{130}$ GFP-Nup49 *lacO*/GFP-LacI | *mus81::KANMX* (background: CFY #2744) | CFY #4746, #4748, #4788<br>This study |
| CFP-LacI, RFP-Nup49 | *MATa, nup49: mCherry-NUP49, ARS607-LacOR-TRP1, ura3-1:: HIS3p-CFP-LacI-URA3, leu2-3, 112 trp1-1 ade2-1 can1-100;Chr6int1::lacop-lexAop-TRP1 (Chr6int1 is 272 bp upstream ARS607)* | CFY #6139<br>This study |
| $(AT)_{34}$ RFP-Nup49 *lacO*/CFP-LacI | *Chr6int2::S5'-Flex1*$(AT)_{34}$*-S3'-HPH* (Chr6int2 is 6371 bp from Chr6int1 and 560bp upstream of Ta(AGC)F gene) (background: CFY #6139) | CFY #6157, #6158<br>This study |
| *Rad51-YFP* $(AT)_{34}$ RFP-Nup49 *lacO*/CFP-LacI | *Rad51-i-mVenus* (internal tag at L21) (background: CFY #6157) | CFY #6184 & #6185<br>This study |
| Rad51 and Rfa3 colocalization assay strains | | |
| No repeat control RFP-Nup49 *lacO*/CFP-LacI | *Chr6int2::No repeat control-HPH* (Chr6int2 is 6371 bp from Chr6int1 and 560bp upstream of Ta(AGC)F gene) (background: CFY #6139) | CFY #6173 & #6174 |

| | | |
|---|---|---|
| *Rad51-YFP* No repeat control RFP-Nup49 *lacO*/CFP-LacI | *Rad51-i-mVenus* (internal tag at L21) (background: CFY #6173) | CFY #6349<br>This study |
| *mus81Δ Rad51-YFP* $(AT)_{34}$ RFP-Nup49 *lacO*/CFP-LacI | *mus81::KANMX* (background: CFY #6184) | CFY #6235 & #6236<br>This study |
| *mre11D56N Rad51-YFP* $(AT)_{34}$ RFP-Nup49 *lacO*/CFP-LacI | Mating #6184 x #6043 | CFY #6289 & #6290<br>This study |
| *mus81Δmre11D56N Rad51-YFP* $(AT)_{34}$ RFP-Nup49 *lacO*/CFP-LacI | *mus81::KANMX* (background: CFY #6289) | CFY #6306 & #6307<br>This study |
| *Rfa3-YFP* No repeat control RFP-Nup49 *lacO*/CFP-LacI | *Rfa3-mvenus-KANMX* (background: CFY #6173) | CFY #6444 & #6445<br>This study |
| *Rfa3-YFP* $(AT)_{34}$ RFP-Nup49 *lacO*/CFP-LacI | *Rfa3-mvenus-KANMX* (background: CFY #6157) | CFY #6442 & #6443<br>This study |
| Okazaki fragment sequencing strains | | |
| *cdc9::tetO7-CDC9* | *MATalpha ade2-1 trp1-1 can1-100 leu2-3,112 his3-11,15 ura3-1 cdc9::tetO7-CDC9 cmv_Laci-NAT* | CFY #4697/DS #12 |
| ChrVI No repeat control | *MAT? ade2-1 trp1-1 can1-100 leu2-3,112 his3-11,15 ura3-1, tor1-1::HIS3, fpr1::NatMX4, RPL13A-2xFKBP12::TRP1 CDC9-FRB::KanMX, NoRepeatControl-HPH cassette at Chr6int2 Locus* | DS #2442 |
| ChrVI *S5'-Flex1(AT)$_{34}$-S3'* | *MAT? ade2-1 trp1-1 can1-100 leu2-3,112 his3-11,15 ura3-1, tor1-1::HIS3, fpr1::NatMX4, RPL13A-2xFKBP12::TRP1 CDC9-FRB::KanMX ,S5'-(AT)$_{34}$-S3'-HPH cassette at Chr6int2 Locus* | DS #2441 |
| *rad52Δ* ChrVI S5'-Flex1$(AT)_{34}$-S3'<br>(Also used for *rad52Δ* ChrX No insert control) | *MAT? ade2-1 trp1-1 can1-100 leu2-3,112 his3-11,15 ura3-1, tor1-1::HIS3, fpr1::NatMX4, RPL13A-2xFKBP12::TRP1 CDC9-FRB::KanMX, rad52::LEU,S5'-* | DS#2439 |

| | | |
|---|---|---|
| | *$(AT)_{34}$-S3'-HPH cassette at Chr6int2 Locus* | |
| *rad52Δ* ChrVI No repeat control (Also used for *rad52Δ* ChrX $(AT)_{34}$) | *MAT? ade2-1 can1-100 his3-11,15 leu2-3,112 trp1-1 ura3-1 RAD5+, S5'-$(AT)_{34}$-S3' 2 kb downstream ARS1019 cdc9::tetO7-CDC9 cmv_Laci-NAT Rad52::KanMX* | DS#2297 |
| ChrX No insert control | *MAT$\alpha$ ade2-1 can1-100 his3-11,15 leu2-3,112 trp1-1 ura3-1 RAD5+, Flex1 Control 2 kb downstream ARS1019, cdc9::tetO7-CDC9 cmv_Laci-NAT* | DS #2092 |
| ChrX S5'-Flex1$(AT)_{34}$-S3' | *MAT? ade2-1 can1-100 his3-11,15 leu2-3,112 trp1-1 ura3-1 RAD5+, S5'-Flex1$(AT)_{34}$-S3' 2 kb downstream ARS1019 cdc9::tetO7-CDC9 cmv_Laci-NAT* | DS#2258 |
| *rad52Δ* ChrX S5'-Flex1$(AT)_{34}$-S3' (Also used for ChrVI No insert *rad52Δ* control) | *MAT? ade2-1 can1-100 his3-11,15 leu2-3,112 trp1-1 ura3-1 RAD5+, S5'-$(AT)_{34}$-S3' 2 kb downstream ARS1019 cdc9::tetO7-CDC9 cmv_Laci-NAT Rad52::KanMX* | DS#2297 |
| Live-cell imaging of replication fork progression | | |
| Live cell imaging no insert | *leu2-3,112 trp1-1 can1-100 ura3-1 ade2-1 his3-11,15 LacI-Envy and TetR-tdTomato replacing ADE1 with KAN marker LacOx128 array integrated at chrIV:332960. TetOx128 array integrated at chrIV:352560. NAT marker integrated at chrIV:340385* | CFY #6212 (Dovrat et al., 2018) |
| No repeat control | *Chr4int1::No repeat control-HPH (Chr4int1 is 340385)* (background: CFY #6212) | CFY #6296 & #6297 This study |
| *mus81Δ* no repeat control | *mus81::NATMX* (background: CFY #6296) | CFY #6390 This study |
| *rev3Δ* no repeat control | *rev3Δ::NATMX* | CFY #6436 & #6437 |

| | | |
|---|---|---|
| | (background: CFY #6296) | This study |
| (AT)$_{8}$ | *Chr4int1::S5'-Flex1(AT)$_{8}$-S3'-HPH (Chr4int1 is 340385)* (background: CFY #6212) | CFY #6298<br>This study |
| (AT)$_{34}$ | *Chr4int1::S5'-Flex1(AT)$_{34}$-S3'-HPH (Chr4int1 is 340385)* (background: CFY #6212) | CFY #6363<br>This study |
| *mus81Δ* (AT)$_{34}$ | *mus81::NATMX* (background: CFY #6363) | CFY #6392 & #6393<br>This study |
| *sae2* (AT)$_{34}$ | *sae2::NATMX* (background: CFY #6363) | CFY #6397<br>This study |
| *rev3Δ* (AT)$_{34}$ | *rev3Δ::NATMX* (background: CFY #6363) | CFY #6431 & #6432<br>This study |

**Table S2:** Zoning assay individual points for Figure 1B

| Strain | Strain No. | No. Zone 1 | % Zone 1 | No. Zone 2 | % Zone 2 | No. Zone 3 | % Zone 3 | Total No. |
|---|---|---|---|---|---|---|---|---|
| WT no repeat Control | 4839 | 71 | 30.5% | 46 | 19.7% | 116 | 49.8% | 233 |
| | 4840 | 39 | 26.0% | 22 | 14.7% | 89 | 59.3% | 150 |
| WT (AT)$_{14}$ | 4754 | 55 | 28.2% | 55 | 28.2% | 85 | 43.6% | 195 |
| | 4755 | 72 | 36.4% | 39 | 19.7% | 87 | 43.9% | 198 |
| WT (AT)$_{23}$ | 5667 | 46 | 46.0% | 9 | 9.0% | 45 | 45.0% | 100 |
| | 5668 | 60 | 34.3% | 23 | 13.1% | 92 | 52.6% | 175 |
| WT (AT)$_{34}$ | 4841 | 60 | 60.6% | 25 | 25.3% | 14 | 14.1% | 99 |
| | 4842 | 57 | 62.6% | 14 | 15.4% | 20 | 22.0% | 91 |

**Table S3:** Zoning assay raw data

| **Strain and Condition** | **Strain No.** | **Zone 1 foci per strain** | | | **Total Zone 1 foci** | | | **p-value (Fisher's exact to WT (AT)$_{34}$ unless otherwise indicated** |
|---|---|---|---|---|---|---|---|---|
| | | **No.** | **%** | **Total No. cells per strain** | **No.** | **%** | **Total No. cells** | |
| Figure 1B | | | | | | | | |
| WT no repeat control | 4839 | 71 | 30.5% | 233 | 110 | 28.7% | 383 | - |
| | 4840 | 39 | 26.0% | 150 | | | | |
| WT (AT)$_{14}$ | 4754 | 55 | 28.2% | 195 | 127 | 32.3% | 393 | =0.3109 vs control |
| | 4755 | 72 | 36.4% | 198 | | | | |
| WT (AT)$_{23}$ | 5067 | 46 | 46.0% | 100 | 106 | 38.5% | 275 | =0.0091 vs control |
| | 5068 | 60 | 34.3% | 175 | | | | |

| | | | | | | | | |
|---|---|---|---|---|---|---|---|---|
| WT $(AT)_{34}$ | 4841 | 60 | 60.6% | 99 | 117 | 61.6% | 190 | <0.0001 vs control |
| | 4842 | 57 | 62.6% | 91 | | | | |
| Figure 1D | | | | | | | | |
| WT $(AT)_{34}$ 30mins | 4841 | 33 | 26.2% | 126 | 84 | 31.2% | 269 | - |
| | 4842 | 51 | 35.7% | 143 | | | | |
| WT $(AT)_{34}$ 30mins + HU | 4841 | 34 | 31.2% | 109 | 60 | 31.3% | 192 | =1.0 vs WT $(AT)_{34}$ 30’ |
| | 4842 | 26 | 31.3% | 83 | | | | |
| WT $(AT)_{34}$ 60mins | 4841 | 72 | 55.8% | 129 | 125 | 53.6% | 233 | - |
| | 4842 | 53 | 51.0% | 104 | | | | |
| WT $(AT)_{34}$ 60mins + HU | 4841 | 31 | 29.0% | 107 | 58 | 33.9% | 171 | =0.0001 vs WT $(AT)_{34}$ 60’ |
| | 4842 | 27 | 42.2% | 64 | | | | |
| WT $(AT)_{34}$ 90mins | 4842 | 41 | 28.5% | 144 | 41 | 28.5% | 144 | |
| WT $(AT)_{34}$ 90mins + HU | 4842 | 81 | 59.1% | 137 | 81 | 59.1% | 137 | =0.0001 vs WT $(AT)_{34}$ 90’ |
| Figure 3A | | | | | | | | |
| WT $(AT)_{34}$ | - | | | | 117 | 61.6% | 190 | - |
| *slx5Δ* $(AT)_{34}$ | 5338 | 31 | 39.2% | 79 | 82 | 40.4% | 203 | <0.0001 |
| | 5339 | 51 | 41.1% | 124 | | | | |
| *uls1Δ* $(AT)_{34}$ | 6512 | 35 | 35% | 100 | 83 | 40.7% | 204 | <0.0001 |
| | 6513 | 48 | 46.2% | 104 | | | | |
| *uls1Δslx5Δ* $(AT)_{34}$ | 6536 | 32 | 30.5% | 105 | 65 | 34.0% | 191 | <0.0001 |
| | 6537 | 33 | 38.4% | 86 | | | | |
| *smt3-331* $(AT)_{34}$ | 5854 | 13 | 26.5% | 49 | 27 | 23.7% | 114 | <0.0001 |
| | 5855 | 14 | 21.5% | 65 | | | | |
| *smt3-KallR* $(AT)_{34}$ | 6489 | 14 | 28.6% | 49 | 50 | 23.3% | 215 | <0.0001 |
| | 6490 | 36 | 21.7% | 166 | | | | |
| *mms21-11* $(AT)_{34}$ | 5487 | 20 | 33.3% | 60 | 39 | 29.8% | 131 | <0.0001 |
| | 5488 | 19 | 26.8% | 71 | | | | |
| *siz1Δ* $(AT)_{34}$ | 5505 | 60 | 50.4% | 119 | 94 | 49.2% | 191 | =0.0178 |
| | 5506 | 34 | 47.2% | 72 | | | | |
| *siz2Δ* $(AT)_{34}$ | 5507 | 57 | 47.5% | 120 | 115 | 45.5% | 253 | =0.0008 |
| | 5508 | 58 | 43.6% | 133 | | | | |
| *siz1Δsiz2Δ* $(AT)_{34}$ | 5588 | 20 | 21.7% | 92 | 29 | 22.0% | 132 | <0.0001 |
| | 5589 | 9 | 22.5% | 40 | | | | |
| Figure 3B | | | | | | | | |
| WT | | | | | 117 | 61.6% | 190 | - |

| | | | | | | | | |
|---|---|---|---|---|---|---|---|---|
| *rad52Δ* (AT)$_{34}$ | 5775 | 92 | 55.1% | 167 | 172 | 58.7% | 293 | =0.5690 |
| | 5776 | 80 | 63.5% | 126 | | | | |
| *dnl4Δ* (AT)$_{34}$ | 6506 | 59 | 50.0% | 118 | 141 | 49.6% | 284 | 0.0112 |
| | 6507 | 82 | 49.4% | 166 | | | | |
| *rad51Δ* (AT)$_{34}$ | 5445 | 47 | 34.3% | 137 | 89 | 36.6% | 243 | <0.0001 |
| | 5446 | 42 | 39.6% | 106 | | | | |
| *rad51-II3A* (AT)$_{34}$ | 5785 | 75 | 36.8% | 204 | 141 | 32.6% | 433 | <0.0001 |
| | 5786 | 66 | 28.8% | 229 | | | | |
| *PCNA-K164R* (AT)$_{34}$ | 6035 | 80 | 56.7% | 141 | 138 | 56.8% | 243 | =0.3266 |
| | 6036 | 58 | 56.9% | 102 | | | | |
| *rad5Δ* (AT)$_{34}$ | 6005 | 24 | 53.3% | 45 | 102 | 65.4% | 156 | =0.5020 |
| | 6006 | 78 | 70.3% | 111 | | | | |
| *rev3Δ (AT)$_{34}$* | 6213 | 47 | 50.0% | 94 | 113 | 50.9% | 222 | =0.0365 |
| | 6248 | 66 | 51.6% | 128 | | | | |
| Figure 2A & Figure S2A | | | | | | | | |
| WT (AT)$_{34}$ | | | | | 117 | 61.6% | 190 | - |
| *yen1Δ* (AT)$_{34}$ | 5150 | 80 | 70.8% | 113 | 137 | 72.1% | 190 | =0.0382 |
| | 5151 | 57 | 74.0% | 77 | | | | |
| *slx1Δ* (AT)$_{34}$ | 6148 | 38 | 33.0% | 115 | 59 | 32.2% | 183 | <0.0001 |
| | 6149 | 21 | 30.9% | 68 | | | | |
| *slx4Δ* (AT)$_{34}$ | 5336 | 62 | 36.0% | 172 | 120 | 30.8% | 390 | <0.0001 |
| | 5931 | 35 | 26.5% | 132 | | | | |
| | 5932 | 23 | 26.7% | 86 | | | | |
| *mus81Δ* (AT)$_{34}$ | 4922 | 19 | 32.8% | 58 | 108 | 34.7% | 311 | <0.0001 |
| | 4923 | 11 | 34.4% | 32 | | | | |
| | 5744 | 46 | 38.7% | 119 | | | | |
| | 5755 | 32 | 31.4% | 102 | | | | |
| *mms4Δ* (AT)$_{34}$ | 5517 | 20 | 38.5% | 52 | 74 | 40.2% | 184 | <0.0001 |
| | 5518 | 54 | 40.9% | 132 | | | | |
| *rad1Δ* (AT)$_{34}$ | 6135 | 39 | 41.1% | 95 | 66 | 42.0% | 157 | =0.0004 |
| | 6136 | 27 | 43.5% | 62 | | | | |
| *sae2Δ* (AT)$_{34}$ | 5597 | 56 | 44.4% | 126 | 90 | 44.6% | 202 | =0.0008 |
| | 5598 | 34 | 44.7% | 76 | | | | |
| *mre11D56N* (AT)$_{34}$ | 6041 | 34 | 36.7% | 93 | 68 | 38.6% | 176 | <0.0001 |
| | 6042 | 34 | 41.0% | 83 | | | | |
| *mus81Δsae2Δ* (AT)$_{34}$ | 5796 | 34 | 25.6% | 133 | 55 | 24.4% | 225 | <0.0001 |
| | 5797 | 21 | 22.8% | 92 | | | | |
| Figure 2B | | | | | | | | |
| WT (CAG)$_{130}$ | 2744 | 78 | 48.2% | 162 | 151 | 47.3% | 319 | - |
| (Maclay et al., 2025) | 3116 | 73 | 46.6% | 157 | | | | |

| | | | | | | | | |
|---|---|---|---|---|---|---|---|---|
| *mus81Δ* $(CAG)_{130}$ | 4746 | 47 | 40.9% | 115 | 164 | 48.7% | 337 | =0.6873 vs WT $(CAG)_{130}$ |
| | 4748 | 74 | 56.9% | 130 | | | | |
| | 4788 | 43 | 46.7% | 92 | | | | |
| Figure 5A & S1A | | | | | | | | |
| WT $(AT)_{34}$ (GFP-LacI/GFP-Nup49) | | | | | 117 | 61.6% | 190 | - |
| WT $(AT)_{34}$ (CFP-LacI/ RFP-Nup49) | 6157 | 56 | 54.9% | 102 | 118 | 58.1% | 203 | =0.5370 vs WT $(AT)_{34}$ |
| | 6158 | 62 | 61.4% | 101 | | | | |
| No repeat control (CFP-LacI/RFP-Nup49) | 6173 | 31 | 21.4% | 145 | 57 | 23.0% | 248 | =0.1170 vs WT no repeat control |
| | 6174 | 26 | 25.2% | 103 | | | | |
| *RAD51-YFP* $(AT)_{34}$ | 6184 | 71 | 61.2% | 116 | 120 | 58.3% | 206 | =0.5386 |
| | 6185 | 49 | 54.4% | 90 | | | | |
| Figure S1B | | | | | | | | |
| WT $(AT)_{34}$ 15mins | 4841 | 24 | 30.8% | 78 | | | | |
| WT $(AT)_{34}$ 25mins | 4841 | 22 | 31.0% | 71 | | | | |
| WT $(AT)_{34}$ 35mins | 4841 | 27 | 39.7% | 68 | | | | |
| WT $(AT)_{34}$ 45mins | 4841 | 41 | 43.5% | 94 | | | | |
| WT $(AT)_{34}$ 55mins | 4841 | 50 | 48.5% | 103 | | | | |
| WT $(AT)_{34}$ 65mins | 4841 | 54 | 54.0% | 100 | | | | |
| WT $(AT)_{34}$ 75mins | 4841 | 49 | 55.1% | 89 | | | | |
| WT $(AT)_{34}$ 85mins | 4841 | 28 | 32.9% | 85 | | | | |

**Table S4:** Colocalization of nuclear periphery with the lacO array 6.4 kb from Flex1$(AT)_{34}$

| **Time after $\alpha$-factor release** | **Live Cell Colocalization WT (AT)34** | | | **Fixed Cell Colocalization WT (AT)34 Exp #1** | | **Fixed Cell Colocalization WT (AT)34 Exp #2** | | **Live Cell Colocalization *yen1Δ* Flex1(AT)34** | | |
|---|---|---|---|---|---|---|---|---|---|---|
| | **% Colocalization** | **No. of Cells colocalized** | **No. Cells** | **% Colocalization** | **No. Cells** | **% Colocalization** | **No. Cells** | **% Colocalization** | **No. of Cells colocalized** | **No. Cells** |
| 10 | 33.3 | 34 | 102 | | | | | 33.3 | 21 | 63 |
| 15 | 38.2 | 42 | 110 | | | 30.4 | 115 | 33.3 | 28 | 84 |
| 20 | 36.9 | 41 | 111 | | | | | 37.5 | 30 | 80 |
| 25 | 35.5 | 38 | 107 | 33.7 | 181 | | | 35.2 | 31 | 88 |
| 30 | 44.6 | 50 | 112 | | | 33.1 | 121 | 32.9 | 25 | 76 |
| 35 | 39.5 | 47 | 119 | | | | | 31.8 | 28 | 88 |
| 40 | 40.0 | 46 | 115 | 36.1 | 144 | | | 40.9 | 36 | 88 |
| 45 | 45.0 | 50 | 111 | | | 41.2 | 114 | 41.9 | 36 | 86 |
| 50 | 39.8 | 45 | 113 | | | | | 44.0 | 40 | 91 |

| | | | | | | | | | | |
|---|---|---|---|---|---|---|---|---|---|---|
| 55 | 42.6 | 46 | 108 | 50.8 | 183 | | | 44.4 | 40 | 90 |
| 60 | 48.1 | 52 | 108 | | | 46.9 | 128 | 53.9 | 48 | 89 |
| 65 | 49.5 | 54 | 109 | | | | | 51.3 | 40 | 78 |
| 70 | 50.9 | 54 | 106 | 52.5 | 162 | | | 55.6 | 45 | 81 |
| 75 | 48.0 | 49 | 102 | | | 55.6 | 117 | 51.9 | 41 | 79 |
| 80 | 47.6 | 49 | 103 | | | | | 46.9 | 38 | 81 |
| 85 | 46.7 | 50 | 107 | 43.9 | 189 | | | 44.1 | 30 | 68 |
| 90 | 50.9 | 52 | 102 | | | 40.7 | 123 | 42.3 | 30 | 71 |
| 95 | 44.4 | 44 | 99 | | | | | 36.8 | 25 | 68 |
| 100 | 38.4 | 38 | 99 | 42.1 | 171 | | | 35.4 | 23 | 65 |

**Table S5:** Rates of deletion measured by the direct duplication recombination assay (DDRA)

| **Strain Name (Strain Number)** | **Individual rate of $FOA^R$ $Ade^+$ ($x10^{-5}$)** | **Average rate of $FOA^R$ $Ade^+$ ($x10^{-5}$) ± SEM** | **Fold-over WT $(AT)_{34}$** | **p-value with respect to WT $(AT)_{34}$ unless otherwise indicated** |
|---|---|---|---|---|
| Figure 1E | | | | |
| WT $(AT)_{34}$ (2525) | 26.91<br>19.54<br>26.91<br>37.66<br>41.01<br>32.82<br>41.92<br>29.80<br>37.60<br>29.76<br>27.44<br>28.78<br>37.57<br>39.70<br>29.50<br>30<br>33 | 32.56 ± 1.406 | - | - |
| WT $(AT)_{34}$ (2712) | 25.99<br>36.14 | | | |
| *nup84Δ* $(AT)_{34}$ (5672) | 106.36<br>75.54 | 74.66 ± 11.97 | 2.3x | =0.0410 |
| *nup84Δ* $(AT)_{34}$ (5673) | 67.91<br>48.82 | | | |
| *mps3Δ75-150* (5634) | 42.25 | 34.60 ± 3.007 | 1x | =0.7965 |

| | | | | |
|---|---|---|---|---|
| *mps3Δ75-150* (5653) | 25.29 | | | |
| *mps3Δ75-150* (5726) | 41.88 | | | |
| *mps3Δ75-150* (5727) | 29.91<br>39.17<br>29.1 | | | |
| *slx5Δ* $(AT)_{34}$ (5711) | 67.04<br>71.07 | 80.74 ± 8.720 | 2.5x | =0.0106 |
| *slx5Δ* $(AT)_{34}$ (5712) | 105.81<br>79.05 | | | |
| Figure 3C | | | | |
| *dnl4Δ* $(AT)_{34}$ (6508) | 48.47<br>41.44<br>37.0<br>32.63<br>41.67 | 43.28 ± 2.792 | 1.3x | =0.0072 |
| *dnl4Δ* $(AT)_{34}$ (6532) | 54.59<br>47.28 | | | |
| *rad51Δ* $(AT)_{34}$ (#4705) | 37.2<br>36.7<br>39.5<br>49.7 | 40.36 ± 1.52 | 1.2x | =0.0012 |
| *rad51Δ* $(AT)_{34}$ (#4708) | 38.2<br>36.2<br>44.2<br>37.6<br>43.9 | | | |
| *rad51-II3A* $(AT)_{34}$ (5783) | 60.96<br>75.53 | 67.25 ±3.6529 | 2x | =0.001 |
| *rad51-II3A* $(AT)_{34}$ (5784) | 61.25<br>71.25 | | | |
| WT no repeat control (2864) (Das et al., 2025) | 2.9<br>1.9<br>3.0<br>2.7<br>2.3<br>3.8<br>3.7<br>3.5<br>2.3 | 2.9 ± 0.2 | - | - |

| | | | | |
|---|---|---|---|---|
| *dnl4Δ* no repeat control (6510) | 3.35<br>1.68<br>3.47 | 3.964 $\pm$ 0.834 | 1.3x | =0.2774<br>vs WT no repeat control |
| *dnl4Δ* no repeat control (6511) | 6.74<br>4.58 | | | |
| *rad51* no repeat control (6077) | 12.7<br>13.13 | 13.07 $\pm$ 1.6703 | 4.2x | =0.0091<br>vs WT no repeat control |
| *rad51* no repeat control (6078) | 17.3<br>9.14 | | | |
| *rad51-II3A* no repeat control (6542) | 9.68<br>12.73<br>9.78 | 8.95 $\pm$ 0.778 | 3.1x | =0.0001<br>Vs WT no repeat control |
| *rad51-II3A* no repeat control (6549) | 6.62<br>6.96<br>8.32<br>8.56 | | | |
| Figure 2C | | | | |
| *rad52Δ* (AT)$_{34}$ (5781) | 2.27<br>1.82 | 2.3 $\pm$ 0.1529 | 0.1x | <0.0001 vs WT<br>=0.1087 vs *mus81Δsae2Δ* |
| *rad52Δ* (AT)$_{34}$ (5782) | 2.52<br>2.4 | | | |
| *mus81Δ* (AT)$_{34}$ (Kaushal et al., 2019) | 15.2<br>12.9<br>13.2 | 13.8 $\pm$ 0.7219 | 0.4x | <0.0001 |
| *sae2Δ* (AT)$_{34}$ (Kaushal et al., 2019) | 5.40<br>7.60<br>6.30<br>11.20 | 7.6 $\pm$ 1.274 | 0.2x | <0.0001 |
| *sae2Δmus81Δ* (AT)$_{34}$ (5787) | 6.25<br>3.34<br>4.39<br>2.49 | 4.118 $\pm$ 0.8101 | 0.1x | <0.0001 vs WT<br>=0.0003 vs *mus81Δ*<br>=0.067 vs *sae2Δ* |

**Table S6**: Rad51 foci counts

| **Strain** | **Strain No.** | **Total counted** | **% of cells containing Rad51 focus/foci** | **No Rad51 focus** | **Rad51 focus** | **Rad51 “rod”** | **p-value** |
|---|---|---|---|---|---|---|---|
| WT no repeat control | #6349 | 406 | 7.9% | 374 | 32 | 0 | - |

| | | | | | | | |
|---|---|---|---|---|---|---|---|
| WT (AT)$_{34}$ | #6184 | 198 | 15.7% | 167 | 31 | 0 | =0.0044 compared to control |
| *mus81Δ* (AT)$_{34}$ | #6235 | 154 | 21.4% | 121 | 29 | 4 | |
| *mre11D56N* (AT)$_{34}$ | #6289 | 155 | 29.0% | 110 | 44 | 1 | =0.0027 compared to WT (AT)$_{34}$ |
| *mus81Δmre11D56N* (AT)$_{34}$ | #6307 | 213 | 20.6% | 169 | 38 | 6 | |

**Table S7**: Colocalization of Rad51 foci with the lacO array 6.4 kb from Flex1(AT)$_{34}$

| Strain | Strain No. | No. Rad51-(AT)$_{34}$ colocalized | % Rad51-(AT)$_{34}$ colocalized | No. Rad51-(AT)$_{34}$ NOT colocalized | Total Rad51 foci counted | % Colocalized per condition | p-value |
|---|---|---|---|---|---|---|---|
| WT no repeat control | #6349 | 2 | 3.8% | 50 | 52 | 3.8% | |
| WT (AT)$_{34}$ | #6184 | 13 | 14.0% | 80 | 93 | 14.9% | =0.0467 vs no repeat control |
| | #6185 | 10 | 16.4% | 51 | 61 | | |
| *mus81Δ* (AT)$_{34}$ | #6235 | 4 | 8.3% | 44 | 48 | 8.0% | =0.1180 vs (AT)$_{34}$ WT |
| | #6236 | 4 | 7.7% | 48 | 52 | | |
| *mre11D56N* (AT)$_{34}$ | #6289 | 32 | 44.4% | 40 | 72 | 44.2% | <0.0001 vs (AT)$_{34}$ WT |
| | #6290 | 29 | 43.9% | 37 | 66 | | |
| *mus81Δmre1156N* (AT)$_{34}$ | #6306 | 5 | 8.8% | 57 | 62 | 9.6% | <0.0001 vs *Mre11D56N* |
| | #6307 | 5 | 11.9% | 37 | 42 | | |

**Table S8**: Nuclear location of Rad51 foci colocalized with the lacO array/Flex1(AT)$_{34}$ locus

| Strain | Strain No. | No. Rad51-(AT)$_{34}$ colocalized at NP | No. Rad51-(AT)$_{34}$ colocalized in nuclear interior | Total Rad51 colocalization events counted | % Rad51-(AT)$_{34}$ colocalized in nuclear interior per strain | % Rad51-(AT)$_{34}$ colocalized in nuclear interior per condition |
|---|---|---|---|---|---|---|
| No repeat control | #6349 | 0 | 2 | 2 | 100% | 100% |
| WT (AT)$_{34}$ | #6184 | 4 | 8 | 12 | 66.7% | 68.2% |
| | #6185 | 3 | 7 | 10 | 70.0% | |
| *mus81Δ* (AT)$_{34}$ | #6235 | 0 | 4 | 4 | 100% | 87.5% |
| | #6236 | 1 | 3 | 4 | 75% | |
| *mre11D56N* (AT)$_{34}$ | #6289 | 5 | 27 | 32 | 84.4% | 80.3% |
| | #6290 | 7 | 22 | 29 | 75.9% | |

| | | | | | | |
|---|---|---|---|---|---|---|
| *mus81Δmre1156N* (AT)$_{34}$ | #6306 | 0 | 5 | 5 | 100% | 100% |
| | #6307 | 0 | 5 | 5 | 100% | |

**Table S9**: Rfa3 foci counts

| **Strain** | **Strain No.** | **Total counted** | **% of cells containing Rfa3 focus/foci** | **Rfa3 focus** | **Combined % of cells containing Rfa3 focus/foci** | **p-value** |
|---|---|---|---|---|---|---|
| WT no repeat control | #6444 | 655 | 22.4% | 147 | 22.4% | - |
| | #6445 | 672 | 22.3% | 150 | | |
| WT (AT)$_{34}$ | #6442 | 450 | 18.9% | 85 | 18.6% | =0.0311 |
| | #6443 | 485 | 18.4% | 89 | | |

**Table S10**: Colocalization of Rfa3 foci with the lacO array 6.4 kb from Flex1(AT)$_{34}$

| **Strain** | **Strain No.** | **No. Rfa3-(AT)$_{34}$ colocalized** | **% Rfa3-(AT)$_{34}$ colocalized** | **No. Rfa3-(AT)$_{34}$ NOT colocalized** | **Total Rfa3 foci counted** | **% Colocalized per condition** | **p-value** |
|---|---|---|---|---|---|---|---|
| WT no repeat control | #6444 | 4 | 5.6% | 71 | 75 | 4.4% | - |
| | #6445 | 2 | 3.1% | 65 | 67 | | |
| WT (AT)$_{34}$ | #6442 | 6 | 5.8% | 98 | 104 | 6.0% | =0.4951 |
| | #6443 | 9 | 6.2% | 136 | 145 | | |

**Table S11**: Nuclear location of Rfa3 foci colocalized with the lacO array/Flex1(AT)$_{34}$ locus

| **Strain** | **Strain No.** | **No. Rad51-(AT)$_{34}$ colocalized at NP** | **No. Rad51-(AT)$_{34}$ colocalized in nuclear interior** | **Total Rad51 colocalization events counted** | **% Rad51-(AT)$_{34}$ colocalized in nuclear interior per strain** | **% Rad51-(AT)$_{34}$ colocalized in nuclear interior per condition** |
|---|---|---|---|---|---|---|
| WT no repeat control | #6444 | 0 | 4 | 4 | 100% | 83.3% |
| | #6445 | 1 | 1 | 2 | 50% | |
| WT (AT)$_{34}$ | #6442 | 0 | 6 | 6 | 100% | 93.3% |
| | #6443 | 1 | 8 | 9 | 88.9% | |
